# Investigating the potential role of intimin/invasin in Aeromonas hydrophila virulence in Labeo rohita: a host-microbe interaction study

**DOI:** 10.1101/2024.11.13.620960

**Authors:** Agradip Bhattacharyya, Goutam Banerjee, Pritam Chattopadhyay

## Abstract

*Aeromonas hydrophila* is a primary bacterial pathogen affecting freshwater fish, including *Labeo rohita* (rohu), leading to significant losses in aquaculture. This study investigates the probable role of intimin/invasin, known virulence factors in several bacterial pathogens, in the pathogenesis of *A. hydrophila* in *L. rohita*. Using an in-silico approach, we explore the genetic and structural features of these proteins to predict their potential function in facilitating infection. We analysed the distribution of invasin and intimin across 53 *A. hydrophila* genomes and examined their physicochemical properties, including molecular weight and stability. Additionally, we evaluated the secondary structures to understand their functional roles in host-pathogen interactions. Using homology-based modelling, we generated 3D (tertiary) structures of invasin and intimin and selected the most suitable models for in silico docking experiments with all eight Rohu (*Labeo rohita*) β-integrins-a crucial step in understanding their interactions with the fish (Rohu) host cells. Due to the unavailability of the crystal structure of Rohu β-integrins, we performed homology-based modelling prior to the docking experiments. Our findings reveal that invasin and intimin are present in only 6 of the 53 *A. hydrophila* strains examined, primarily in highly virulent strains previously reported. Notably, invasin lacks disulfide bonds and beta turns. 3D modelling indicates a significant binding affinity of invasin with all human β-integrins, suggesting a critical role in host-pathogen interactions.

## 1. Introduction

Bacterial diseases pose one of the most challenging threats to the lives of fish, both in the wild and in captivity. Diseases can spread rapidly through water, often leading to severe health issues for fish populations. In this direction, *Aeromonas hydrophila* is recognized as a versatile fish pathogen that can have catastrophic consequences for the aquaculture industry [1]. Infections in fish can manifest primarily as ulcerative dermatitis [2] septicaemia [3] necrotizing fasciitis [4] piknosis and necrotic damage in spleen and kidney [5]. *A. hydrophila* is responsible for various conditions, including gill degeneration in *Oreochromis niloticus* [6], hepato-pancreatic infection in channel catfish [7], bilateral exophthalmia in Pangasius [8], visceral haemorrhagic septicaemia in cultured Chinese sturgeon [9], and many more. Additionally, *A. hydrophila* is associated with Aeromonad red sore disease and epizootic ulcerative syndrome in carp and other fish[10] underscoring its versatility as a pathogen in aquatic environments.

Autotransporter-3 (AT-3) family of adhesins are the outer membrane (OM) proteins known as Intimin/Invasin (Int/Inv) and are present in pathogenic *E. coli* (Int), *Yersinia* spp. (Inv), and other proteobacteria strains [11]. Invasins are monomeric autotransporter (Type 3) [12] and promotes pathogen translocation on target cell [13]. Furthermore, invasin of *Y. pseudotuberculosis* binds on β1-integrin superfamily proteins on the surface of eukaryotic host cells and trigger the rearrangement of the host cell cytoskeleton and internalization of the bacteria [14]. While, in *Salmonella enterica* invasin promotes bacterial adhesion or invasion on/to host cells [15]. Similarly, another bacterial outer membrane adhesin called Intimin promotes *Escherichia coli* adhesion to intestinal villi in calf and production of AE lesions [16]. Just like invasins, intimins are also monomeric autotransporters [12] which promote adhesion or invasion of *Salmonella enterica* on/to vertebrate host cells [15]. According to Fairman et al., (2012), both invasins and intimins act as central virulence factors in attaching and effacing lesions (AE lesions), as they can bind to host cells through their C-terminal extracellular domains, while their N-terminal β-domains are embedded in the outer bacterial membrane [17].

On the other hand, integrins (αβ-heterodimeric integral proteins) belong to the cell adhesion receptor superfamily of the eukaryotic hosts and are recognized to bind proteins containing the specific sequence Arg-Gly-Asp or RGD [18]. Fish, as vertebrate have eight different β-integrins [19]. Ligand recognition by integrins is primarily mediated by a cationic binding site on the β-integrin, adjacent to the exposed α-integrin [20]. Bacteria have been shown to interact, either directly or indirectly, with integrins, facilitated by a surface-localized ligand encoded by the pathogen [21]. Previous research reported that, several pathogenic bacteria like *Shigella* and *Yersinia* bind to integrin β1 on host cells, facilitating invasion and infection, [22, 23]. It has also been reported that ligands of some fish integrins contain negatively charged amino acids (Asp or Glu) that are directly involved in receptor binding [24]. Additionally, integrin overexpression has been linked to increased bacterial adhesion and infection in blackwater teleosts [25].

Despite the significance of *A. hydrophila* as a fish pathogen, there is limited information on its autotransporter type-3 proteins, invasin and intimin, and their interactions with fish during disease establishment. In this investigation, we studied the distribution of invasin and intimin across all available *A. hydrophila* genomes. Since tertiary protein structures for *A. hydrophila* invasin/intimin and Rohu β-integrins were unavailable, homology modeling was conducted. Finally, the interactions between invasin/intimin and fish β-integrins were elucidated through in-silico docking experiments.

## 2. Material and Methods

### 2.1. Data acquisition from different databases

The nucleotide sequences for two key test proteins, Invasin (AHA_1064) and Intimin (AHA_1066), were retrieved from the KEGG Orthology database (https://www.genome.jp/kegg/ko.html) (Table S1). These proteins are essential for understanding the pathogenic mechanisms in *Aeromonas hydrophila*. The KEGG Orthology database was chosen for its comprehensive collection of orthologous genes across multiple species, allowing us to accurately identify and retrieve the corresponding gene sequences for these proteins. For functional and structural analysis, the corresponding protein sequences were also necessary. To this end, the protein sequences of Invasin and Intimin, along with the β-integrins (β1-β8) from *Labeo rohita* (Rohu), were obtained from the UniProt database (https://www.uniprot.org/), a leading repository of protein information that includes high-quality, manually curated protein sequences. In addition, the *A. hydrophila* genome sequences were sourced from the NCBI Genome database (https://www.ncbi.nlm.nih.gov/genome), a well-established resource for complete genome sequences. All datasets were accessed and collected on or before March 31, 2024, ensuring that the latest available sequence information was included in the analyses.

### 2.2. Distribution of test proteins within A. hydrophila genomes

We conducted microbial genome BLAST searches using the tBLASTn algorithm (Table S2) to investigate the distribution of the Invasin (AHA_1064) and Intimin (AHA_1066) proteins across *A. hydrophila* genomes. The tBLASTn tool, which compares a protein sequence (query) against nucleotide sequences from microbial genomes, allowed us to explore the presence and conservation of these proteins across various strains of *A. hydrophila*. To further explore synteny (the conserved order of genes) between the invasin and intimin genes and their neighbouring regions in different *A. hydrophila* genomes, we used the SyntTax server. The SyntTax tool allows for the comparison of gene arrangements across bacterial genomes, providing insights into the conservation of genomic contexts. All analyses were performed using online tools and databases accessed on or before March 31, 2024

### 2.3. Prediction of physicochemical properties

The physicochemical properties of the test proteins (Table 1) and β-integrins (Table 2), including amino acid composition, molecular weight, extinction coefficient, aliphatic index, instability index, grand average hydropathy, isoelectric point, molecular formula, total number of atoms, and charged residues, were predicted using the ExPASy-ProtParam server (https://web.expasy.org/protparam/). By predicting these properties for both the test proteins and β-integrins of *Labeo rohita* (Rohu), we were able to gather comprehensive information on the structural and functional attributes of these proteins. This information serves as a foundational dataset for further experimental validations and for understanding the roles these proteins may play in pathogen-host interactions.

**Table 1:** Physicochemical properties of the test proteins selected for this study.

| Physicochemical parameters | Invasin | Intimin |
| --- | --- | --- |
| Number of amino acids | 916 | 535 |
| Molecular weight | 99614.99 | 56335.50 |
| Extinction coefficient (EC) | 127325 M-1cm-1 | 54110 M-1cm-1 |
| Aliphatic Index (AI) | 83.08 | 85.83 |
| Instability index (II) | 29.18 | 25.02 |
| GRAVY (Grand average of hydropathicity) | -0.327 | -0.106 |
| Isoelectric Point (Pi) | 4.77 | 4.42 |
| Molecular Formula | C<sub>4399</sub>H<sub>6864</sub>N<sub>1218</sub>O<sub>1399</sub>S<sub>13</sub> | C<sub>2455</sub>H<sub>3883</sub>N<sub>671</sub>O<sub>824</sub>S<sub>11</sub> |
| Total number of atom | 13893 | 7844 |
| Number of Positively Charged residue<br>(Arg + Lys) | 73 | 30 |
| Number of Negatively Charged residue<br>(Asp + Glu) | 116 | 59 |

**Table 2:** Physicochemical properties of the Rohu β-Integrins.

| Proteins selected for this study | Number of amino acids | Molecular weight | Extinction coefficient (EC) | Aliphatic Index (AI) | Instability index (II) | GRAVY (Grand average of hydropathicity) | Isoelectric Point (Pi) | Molecular Formula | Total number of atom | Number of positively charged residue (Arg + Lys) | Number of Negatively charged residue (Asp + Glu) |
| --- | --- | --- | --- | --- | --- | --- | --- | --- | --- | --- | --- |
| **β1** | 801 | 88914.58 | 62415 | 73.87 | 43.11 | -0.410 | 6.56 | C<sub>3839</sub>H<sub>6119</sub><br>N<sub>1079</sub>O<sub>1203</sub><br>S<sub>71</sub> | 12311 | 98 | 101 |
| **β2** | 767 | 84690.43 | 66425 | 70.90 | 32.88 | -0.410 | 6.13 | C<sub>3649</sub>H<sub>5778</sub><br>N<sub>1030</sub>O<sub>1148</sub><br>S<sub>70</sub> | 11675 | 89 | 96 |
| **β3** | 783 | 85889.65 | 80740 | 77.52 | 44.62 | -0.219 | 5.25 | C<sub>3776</sub>H<sub>5954</sub><br>N<sub>1056</sub>O<sub>1193</sub><br>S<sub>68</sub> | 11845 | 77 | 100 |
| **β4** | 1930 | 214581.74 | 201340 | 73.87 | 43.08 | -0.460 | 5.77 | C<sub>9376</sub>H<sub>14797</sub><br>N<sub>2605</sub>O<sub>2946</sub><br>S<sub>107</sub> | 29831 | 211 | 237 |
| **β5** | 805 | 89154.62 | 69865 | 73.53 | 46.39 | -0.275 | 6.28 | C<sub>3854</sub>H<sub>6071</sub><br>N<sub>1089</sub>O<sub>1195</sub><br>S<sub>74</sub> | 12283 | 88 | 96 |
| **β6** | 779 | 85424.44 | 64810 | 78.31 | 45.60 | -0.105 | 5.05 | C<sub>3711</sub>H<sub>5817</sub><br>N<sub>1007</sub>O<sub>1159</sub><br>S<sub>73</sub> | 11767 | 97 | 68 |
| **β7** | 768 | 85263.04 | 65145 | 77.41 | 39.63 | -0.347 | 6.09 | C<sub>3680</sub>H<sub>5846</sub><br>N<sub>1036</sub>O<sub>1161</sub><br>S<sub>65</sub> | 11788 | 86 | 96 |
| **β8** | 816 | 90464.26 | 85655 | 72.30 | 42.13 | -0.355 | 5.63 | C<sub>3897</sub>H<sub>6102</sub><br>N<sub>1124</sub>O<sub>1222</sub><br>S<sub>69</sub> | 12414 | 81 | 104 |

### 2.4. Secondary structure characterization

The CYS_REC tool (http://linux1.softberry.com) was used to predict disulfide bonds, essential for protein stability and function, including their bonding patterns and the number of cysteine residues involved. Additionally, the SOPMA tool (https://npsa.lyon.inserm.fr/cgibin/npsa_automat.pl?page=/NPSA/npsa_sopma.html) was used to predict the secondary structures of the test proteins, such as alpha helices, extended strands, beta-turns, and random coils. The detail information regarding secondary structure characterizations of the test proteins and β-integrins were tabulated in Table S3 and Table S4, respectively.

### 2.5. Functional analysis

The properties of the test proteins, Invasin and Intimin, were examined using the SOSUI server (http://harrier.nagahama-i-bio.ac.jp/sosui/sosui_submit.html)), which predicts whether a protein is a transmembrane protein based on its amino acid sequence. This analysis provided insights into the membrane-spanning regions of these proteins, crucial for understanding their role in bacterial pathogenesis. Additionally, the CYS_REC program (http://linux1.softberry.com) was employed to estimate the presence or absence of disulfide bonds, which are essential for maintaining the structural stability and proper function of proteins. Disulfide bonds contribute significantly to the folding and stability of extracellular proteins, making them vital for the integrity of proteins involved in host-pathogen interactions. The program also provided detailed analysis of the bonding configuration, and the number of cysteine residues involved in forming these bonds, giving us a better understanding of the structural intricacies of the test proteins. Furthermore, functional analyses were performed for both the test proteins (Table S5) and β-integrins (Table S6), which included evaluating their potential roles in cellular adhesion, signalling, and interactions with other molecules.

### 2.6. Homology modelling and evaluation of the tertiary structure

The 3D structure of the test proteins was predicted using the SWISS-MODEL server, an automated homology modelling tool that leverages templates from the Protein Data Bank (PDB). Structural templates were identified using the ExPASy web server (https://swissmodel.expasy.org/), and the target sequences were aligned with these templates. The SWISS-MODEL server then constructed the models and performed an initial assessment of their quality. To ensure the accuracy and reliability of the predicted 3D structures, the models underwent further evaluation using several analytical tools, including: (a) RAMPAGE [26] (which analyses the quality of the backbone dihedral angles in the protein model, (b) ProQ web-server (https://proq.bioinfo.se/cgi-bin/ProQ/ProQ.cgi), which estimates the quality of the protein model based on its structural features, (c) ProSA-web server (https://prosa.services. came.sbg.ac.at/prosa.php), which assesses the model’s overall quality and identifies potential errors, and (d) Ramachandran plot [27] which evaluates the distribution of dihedral angles to ensure proper protein folding. The tertiary structure of the test proteins (Table S7) and rohu β-Integrins (Table S8) was also evaluated.

### 2.7. Protein-protein Docking

We used the GRAMM and HDOC servers to investigate protein-protein docking interactions. GRAMM (https://gramm.compbio.ku.edu/) predicts various docking poses, including stable and transient interactions, by mapping the intermolecular energy landscape. Whereas, HDOC (http://hdock.phys.hust.edu.cn/) employs a hybrid approach combining template-based modeling and ab initio free docking for protein-protein and protein-DNA/RNA interactions. HDOC accepts amino acid sequences as input and integrates experimental data, such as protein-binding sites and small-angle X-ray scattering information, into the docking process.

## 3. Results

### 3.1. Distribution of invasin and intimin within A. hydrophila strains

In our genomic analysis utilizing the tBLASTn algorithm, we employed protein query sequences from Invasin and Intimin (Table S1) to perform a BLAST search against a comprehensive database of *A. hydrophila* strains. This analysis revealed that Invasin and Intimin were present in only six out of 53 A. hydrophila strains: ATCC 7966, GSH8-2, LP0103, WCX23, WP8-S18-ESBL-02, and 23-C-23 (Table S2). To further investigate the genomic context of these proteins, we conducted a synteny analysis (Fig. 1) which depicts the syntenic relationships of Invasin and Intimin among the six identified strains. The Figure 1 highlighted the conserved regions and any potential rearrangements of genes that may have occurred. This information is critical for understanding the genetic and evolutionary dynamics of *A. hydrophila*, particularly regarding how these virulence factors may influence the pathogenic potential of different strains.

**Fig. 1.**
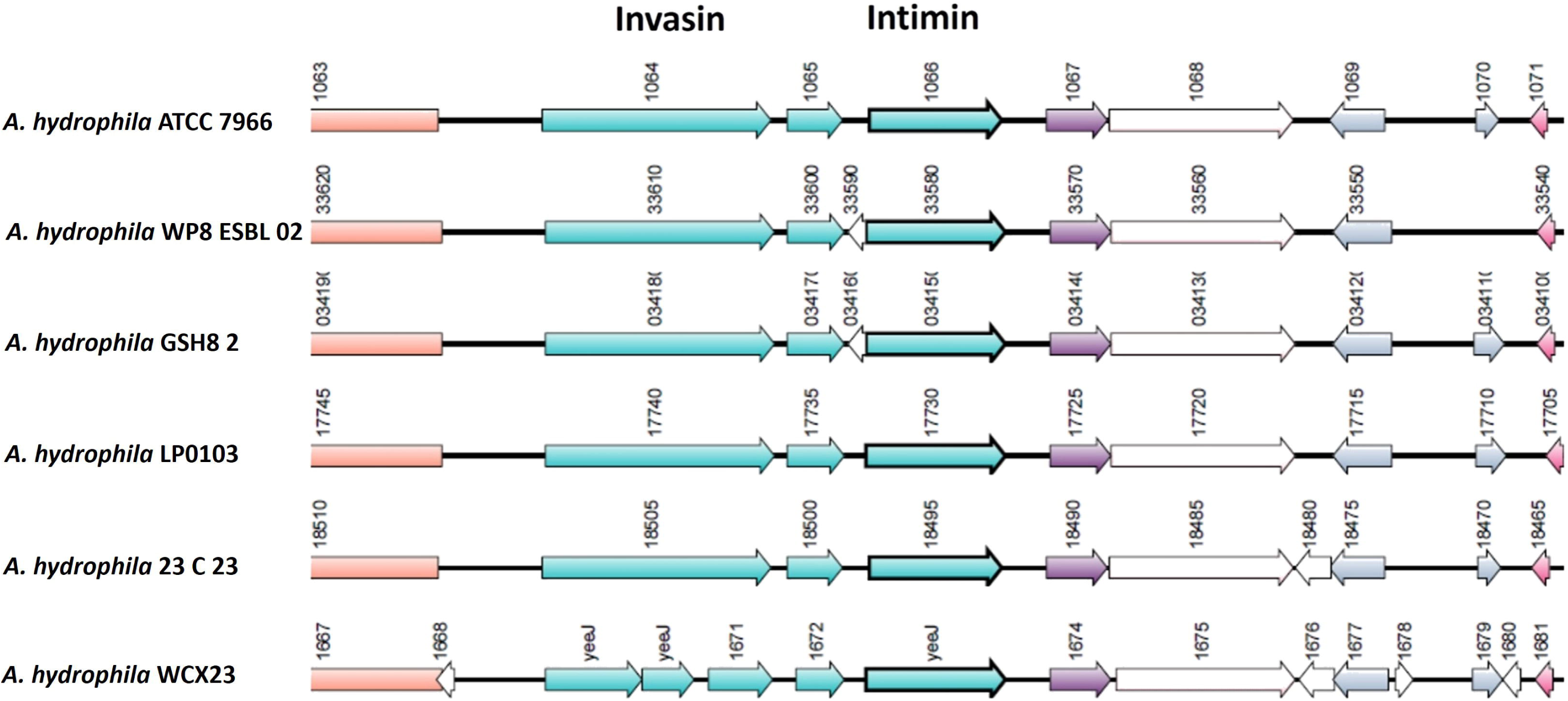
Synteny of invasin and intimin within the *A. hydrophila* genomes.

### 3.2. Physicochemical properties of invasion, intimin and rohu β-Integrins

At 280 nm, the EC (Enzyme commission) of the proteins invasin and intimin were found to be 12732 and 54110 M/cm, respectively (Table 1). Invasin and intimin comprised 916 and 535 amino acids respectively (Table 1). In the present investigation the three experimental proteins may by arranged by pI value in the following manner: Invasin>Intimin. The total amount of negative charge residues (Asp + Glu) in proteins invasin and intimin are 116 and 59 (Table 1). The total quantity of positive charge residues (Arg + Lys) in each of these proteins: invasin and intimin are 73 and 30, respectively. The invasion and intimin exhibit respective instability indexes of 29.18 and 25.02 (Table 1). The aliphatic index (AI) of a protein is defined as the relative volume occupied by aliphatic side chains (alanine, valine, isoleucine, and leucine). It was estimated that the AI for invasin would be 83.08 and intimin would be 85.83 (Table 1).

At 280 nm, the EC of the Rohu β-Integrins (β1 – β8) found to be 62415, 66425, 80740, 201340, 69865, 64810, 65145 and 85655 M/cm, respectively (Table 2). β-Integrins comprised 801, 767,783, 1930, 805, 779, 768 and 816 amino acids respectively (Table 2). In the present investigation the eight Rohu β-Integrins may by arranged by pI value in the following manner: β1 > β5> β2> β7 > β4 > β8 > β3> β6. The total amount of negative charge residues (Asp + Glu) is: 101, 96, 100, 237, 96, 68, 96 and 104 (Table 2). The total quantity of positive charge residues (Arg + Lys) in each of these β-Integrins are 98, 89, 77, 211, 88, 97, 86 and 81. Whereas, Instability Indexes of these β-Integrins are 43.11, 32.88, 44.62, 43.08, 46.39, 45.60, 39.63 and 42.13, respectively (Table 2). The AI for β1, β2, β3, β4, β5, β6, β7 and β8 would be 73.87, 70.90, 77.52, 73.87, 73.53, 78.31, 77.41 and 72.30 respectively (Table 3).

**Table 3:**
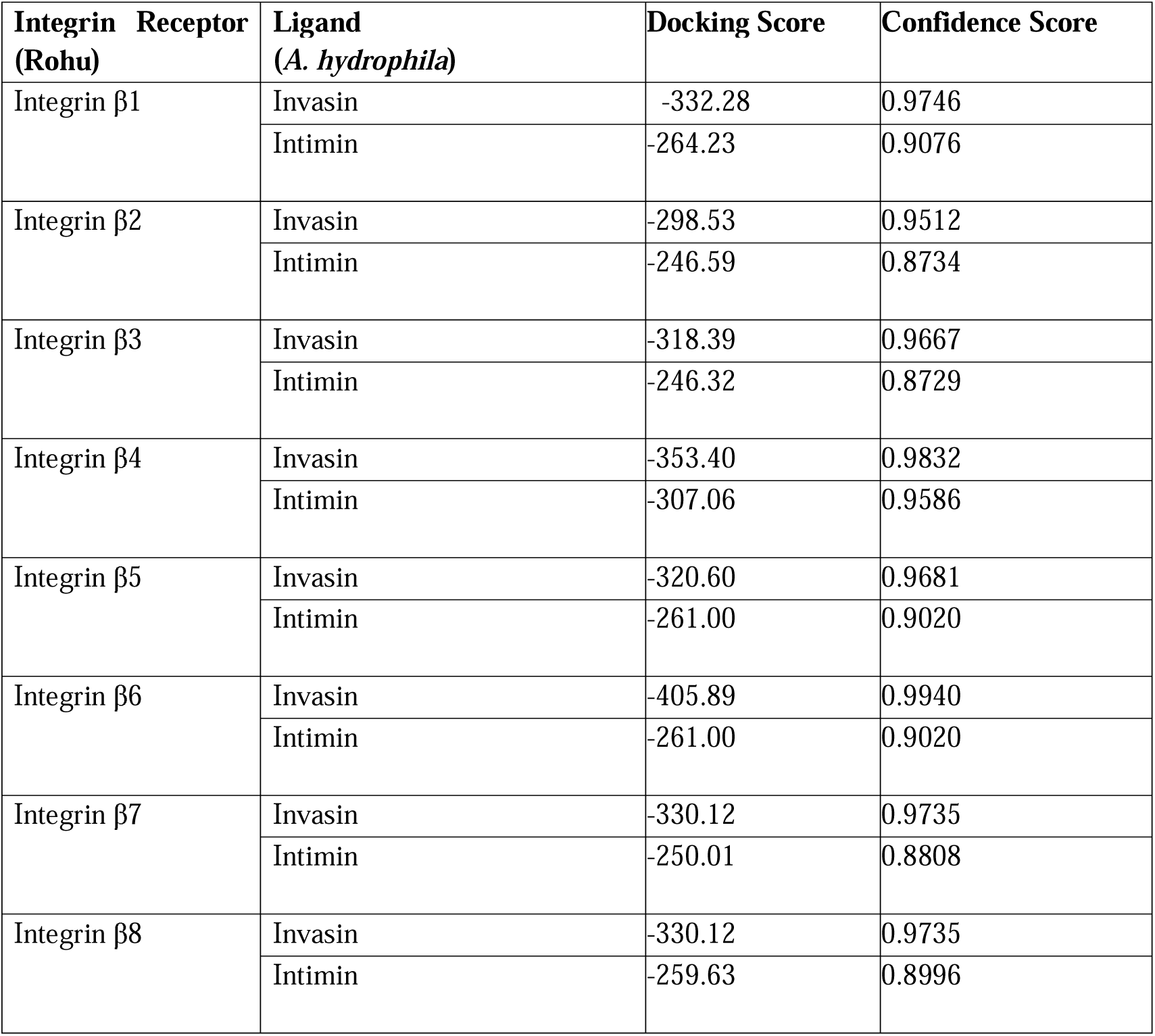
Results of Docking of the test proteins (invasin and intimin) with eight different Human β Integrins.

| **Integrin Receptor**<br>**(Rohu)** | **Ligand**<br>**(*A. hydrophila*)** | **Docking Score** | **Confidence Score** |
| --- | --- | --- | --- |
| Integrin β1 | Invasin | -332.28 | 0.9746 |
|  | Intimin | -264.23 | 0.9076 |
| Integrin β2 | Invasin | -298.53 | 0.9512 |
|  | Intimin | -246.59 | 0.8734 |
| Integrin β3 | Invasin | -318.39 | 0.9667 |
|  | Intimin | -246.32 | 0.8729 |
| Integrin β4 | Invasin | -353.40 | 0.9832 |
|  | Intimin | -307.06 | 0.9586 |
| Integrin β5 | Invasin | -320.60 | 0.9681 |
|  | Intimin | -261.00 | 0.9020 |
| Integrin β6 | Invasin | -405.89 | 0.9940 |
|  | Intimin | -261.00 | 0.9020 |
| Integrin β7 | Invasin | -330.12 | 0.9735 |
|  | Intimin | -250.01 | 0.8808 |
| Integrin β8 | Invasin | -330.12 | 0.9735 |
|  | Intimin | -259.63 | 0.8996 |

### 3.3. Functional analysis of invasion, intimin and rohu β-Integrins

When two cysteines are bonded by an S–S bond, the resulting molecule between the two protein chains is called cystine. Intimin lacks di-sulphide bond (Table S5). However, only two cysteines and single di-sulphide bond was present in the invasin of *Aeromonas hydrophila* (Table S5). On the other hand, 58 Cysteines and 24 di-sulphide bonds, 59 Cysteines and 22 di-sulphide bonds, 56 Cysteines and 32 di-sulphide bonds, 64 Cysteines and 22 di-sulphide bonds, 59 Cysteines and 26 di-sulphide bonds, 57 Cysteines and 25 di-sulphide bonds, 55 Cysteines and 26 di-sulphide bonds and 55 Cysteines and 22 di-sulphide bonds were present in *Labeo rohita* β-Integrins (β1 to β8) respectively (Table S6).

### 3.4. Secondary structure characterization of invasion, intimin and Rohu β-integrins

The secondary structure analysis revealed that the alpha helix constituted 9.72% of the *A. hydrophila* Invasin protein, while Intimin had a significantly lower alpha helix content of only 0.19% (Table S3). In terms of extended strand compositions, Invasin and Intimin displayed values of 25.66% and 29.35%, respectively. Furthermore, approximately 64.63% of Invasin and 70.47% of Intimin were classified as random coils. Notably, neither of these proteins exhibited any beta turns, as illustrated in Figure S1.

In contrast, the analysis of the β-Integrins from *Labeo rohita* (Rohu) demonstrated greater variability in secondary structure composition across the different integrins (β1 to β8). The alpha helix percentages for these integrins were as follows: β1: 20.47%, β2: 20.99%, β3: 21.46%, β4: 11.40%, β5: 20.37%, β6: 21.69%, β7: 22.27%, β8: 23.16% (Table S4). The extended strand compositions for the integrins were β1: 14.73%, β2: 15.12%, β3: 15.96%, β4: 23.99%, β5: 16.27%, β6: 17.46%, β7: 16.02%, β8: 14.46% (Table S4). In terms of random coil content, the integrins exhibited the following percentages: β1: 64.79%, β2: 63.89%, β3: 62.58%, β4: 64.61%, β5: 63.35%, β6: 60.85%, β7: 61.72%, β8: 63.38% (Fig. S2).

### 3.5. Homology modelling of invasin, intimin and Rohu β-integrins

For the Invasin and Intimin proteins from *A. hydrophila* ATCC 7966, 3D models were generated using the SWISS-MODEL server (Table S7). The models were built using the following templates: A0A3T1A166.1.A for invasin (Fig. 2A, GMQE score 0.72), and A0A4P7IWM0.1.A for intimin (Fig. 2B, GMQE score 0.91). The transmembrane regions of these outer membrane proteins were modeled as β-barrels, a common feature of porins (Fig. 2A). The NH2-terminal domains (D1-D4) of Invasin resemble eukaryotic immunoglobulin superfamily (IgSF) folds but lack the disulfide bonds and conserved core residues (Fig. 2B). The D5 domain of Invasin shares structural similarity with C-type lectin-like domains (CTLDs), though it lacks calcium-binding loops. It is composed of two antiparallel β-sheets with α-helical and loop regions and contains a disulfide bond linking helix 1 to β-strand 5, contributing to structural stability (Fig. 2B). These models suggest structural adaptations of Invasin and Intimin, potentially related to their roles in host-pathogen interactions.

**Fig. 2.**
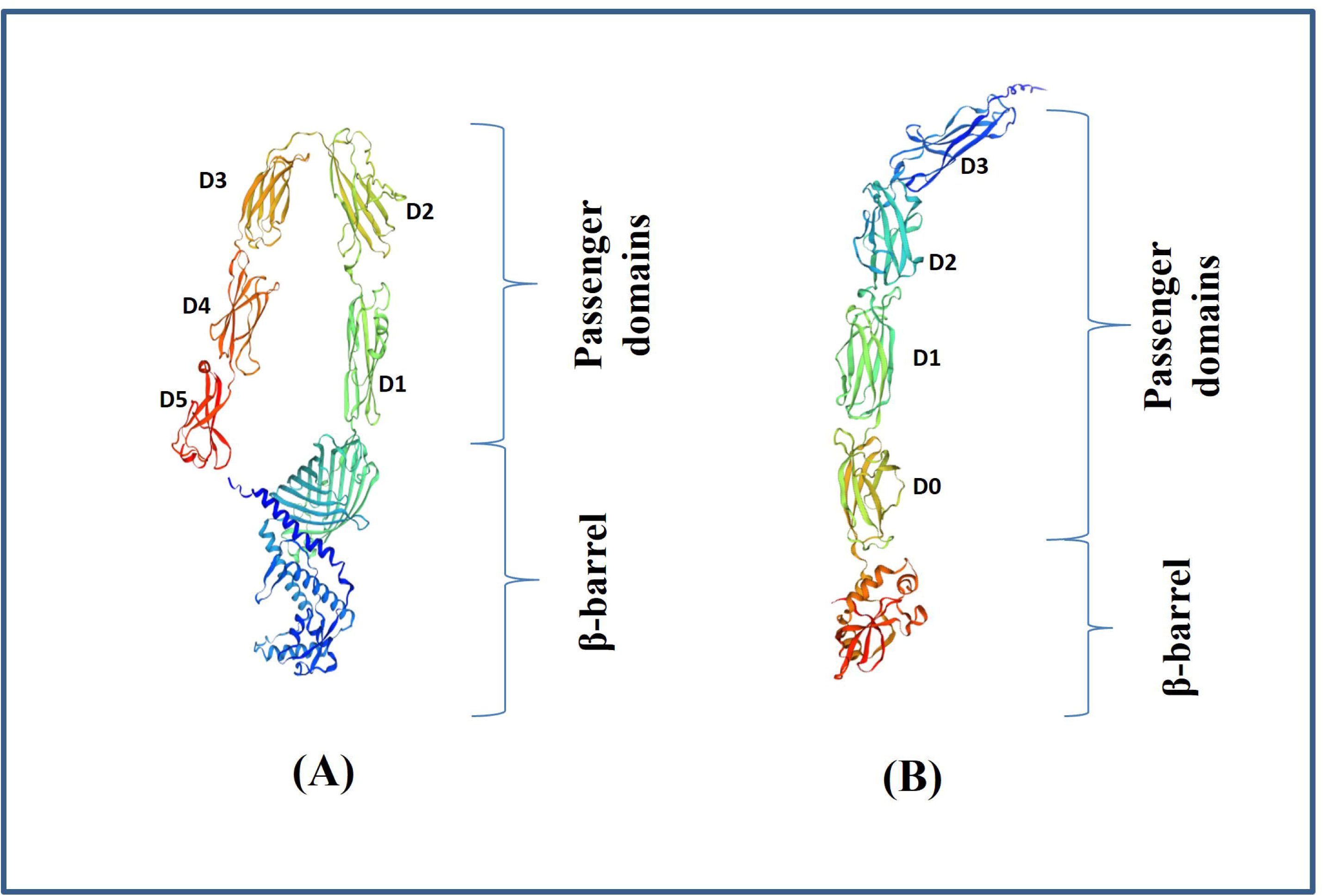
Homology modelling of the (A) invasin and (B) intimin of *A. hydrophila* ATCC 7966 by SWISS-MODEL, accessed via the ExPASy web server.

The most similar 3D models of β-Integrins (β1 to β8) from Rohu were generated using the SWISS-MODEL server based on the best matching sequences (Table S8). The following templates were used for model construction: β1: A0A286Y9W0.1.A (GMQE score: 0.87) (Fig. S3a), β2: E7FCN5.1.A (GMQE score: 0.86) (Fig. S3b), β3: B3DIP9.1.A (GMQE score: 0.84) (Fig. S3c), β4: F1RA51.1.A (GMQE score: 0.71) (Fig. S3d), β5: F1QF91.1.A (GMQE score: 0.83) (Fig. S3e), β6: F1QGX0.1.A (GMQE score: 0.84) (Fig. S3f), β7: E7F4H9.1.A (GMQE score: 0.84) (Fig. S3g), β8: A0A7J6C0U9.1.A (GMQE score: 0.73) (Fig. S3h).

### 3.6. Evaluation of tertiary structure of invasin, intimin & Rohu β-integrins

The structural validation and quality assessment of models for invasin, and intimin were further analysed (Table S9). The results of the ProSAweb validation on the predicted models are represented in three formats (Fig. 3). In Ramachandran plot analysis, the residues in the favoured region of invasin and intimin proteins of *A. hydrophila* were 94.31% and 95.16%, respectively. ProQ server protein model quality analysis revealed that predicted LG score scores of invasin and intimin of *A. hydrophila* were 11.489 and 10.667, respectively. The MaxSub values were –0.871 for Invasin and –1.691 for Intimin, further supporting the quality of the predicted structures.

**Fig. 3.**
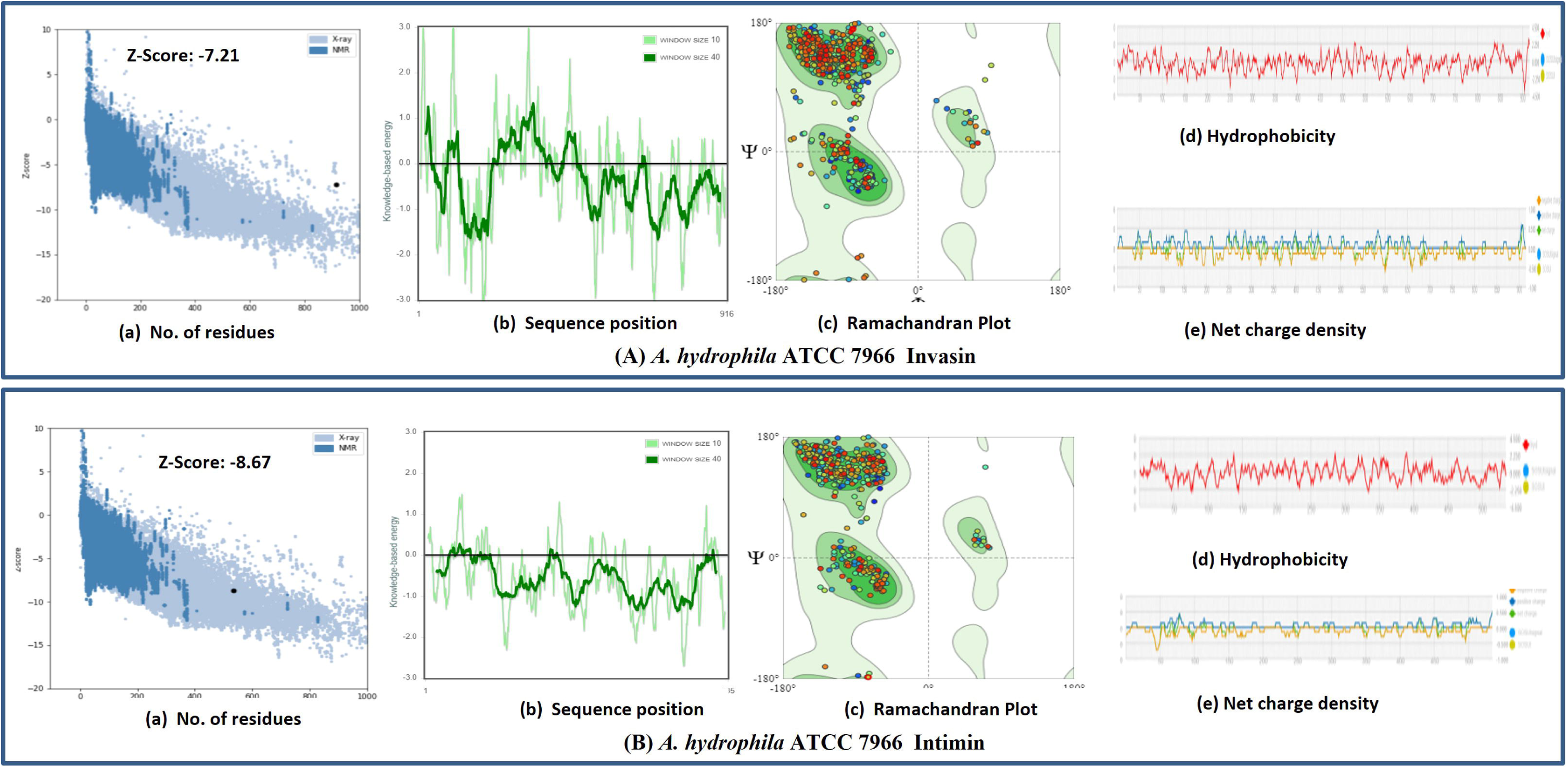
Evaluation of tertiary structures of the (A) invasin and (B) intimin of *A. hydrophila* ATCC 7966. In each panel, (a) represents Z-scores in terms of recediues, (b) represents sequence position, (c) represents Ramachandran plot, (d) represents hydrophobicity, and (e) represents net charge density.

Similarly, the structural validation and quality assessment of the predicted Rohu β-Integrins (β1-β8) models were conducted (Table S10). Results from ProSA-web validation is presented in three formats (Fig. S4). In the Ramachandran plot analysis, the residues in the favoured regions for β-Integrins (β1-β8) were 96.25%, 96.60%, 96.07%, 87.71%, 94.27%, 95.11%, 94.46%, and 92.80%, respectively. The ProQ server revealed LG scores of 9.647, 11.038, 11.218, –0.835, 10.442, 9.884, 10.113, and 9.409, respectively. The MaxSub values were –0.548, –0.659, –0.751, –0.113, –0.594, –0.517, –0.599, and –0.463 for β1-β8, respectively.

### 3.7. Molecular docking of invasin and intimin with Rohu integrins

Key Invasin residues involved in integrin binding include 903-913, forming helix 1 and its adjacent loop in domain D5. The best docking models were selected based on the interactions between integrin, intimin, and Rohu β-integrins, using docking scores (Fig. 4). Models involving the transmembrane domain of Rohu β-integrins were rejected. Docking quality was evaluated through docking score, confidence score (Table 3), and ligand RMSDs (Table S11). Docking scores ranged from –405.89 to –246.23, while confidence scores varied from 0.8734 to 0.9832. RMSDs of residue pairs within 5.0 Å between the receptor and ligand are detailed in Table S11. For instance, Thr 894 of Invasin (D5) interacts with Gln 626 of Integrin β14 (Fig. 5A), and Tyr 39 of Intimin (D3) interacts with Met 689 of Integrin β1 (Fig. 5B). Full residue interactions and RMSD values for all docking models are provided in Table S11.

**Fig. 4.**
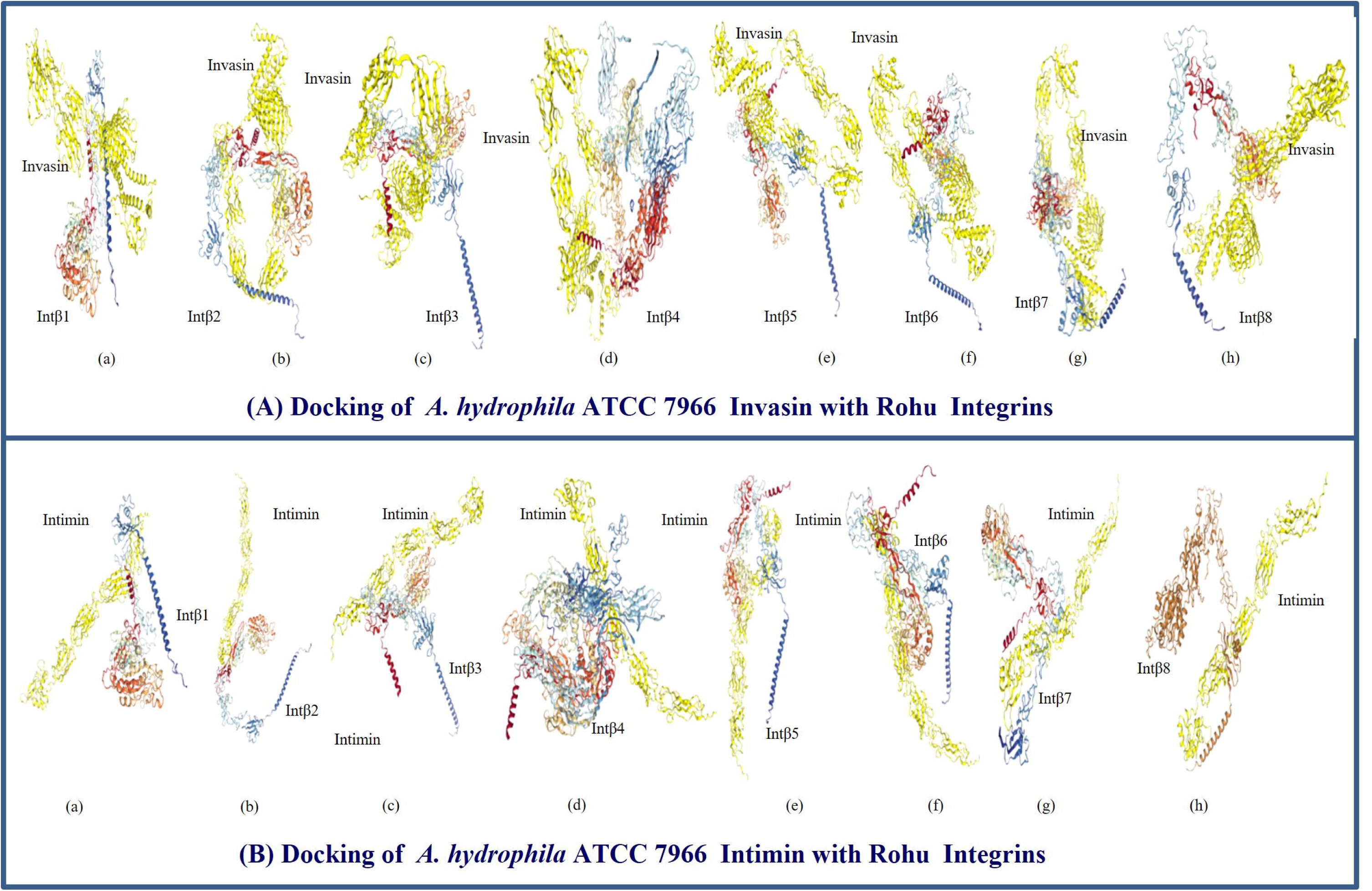
Molecular docking of the (A) invasin and (B) intimin proteins of *A. hydrophila* ATCC 7966 with β-integrins of *L. rohita*. In each panel (A and B), a to h indicates docking of a particular test protein with integrin β1 to β8.

**Fig. 5.**
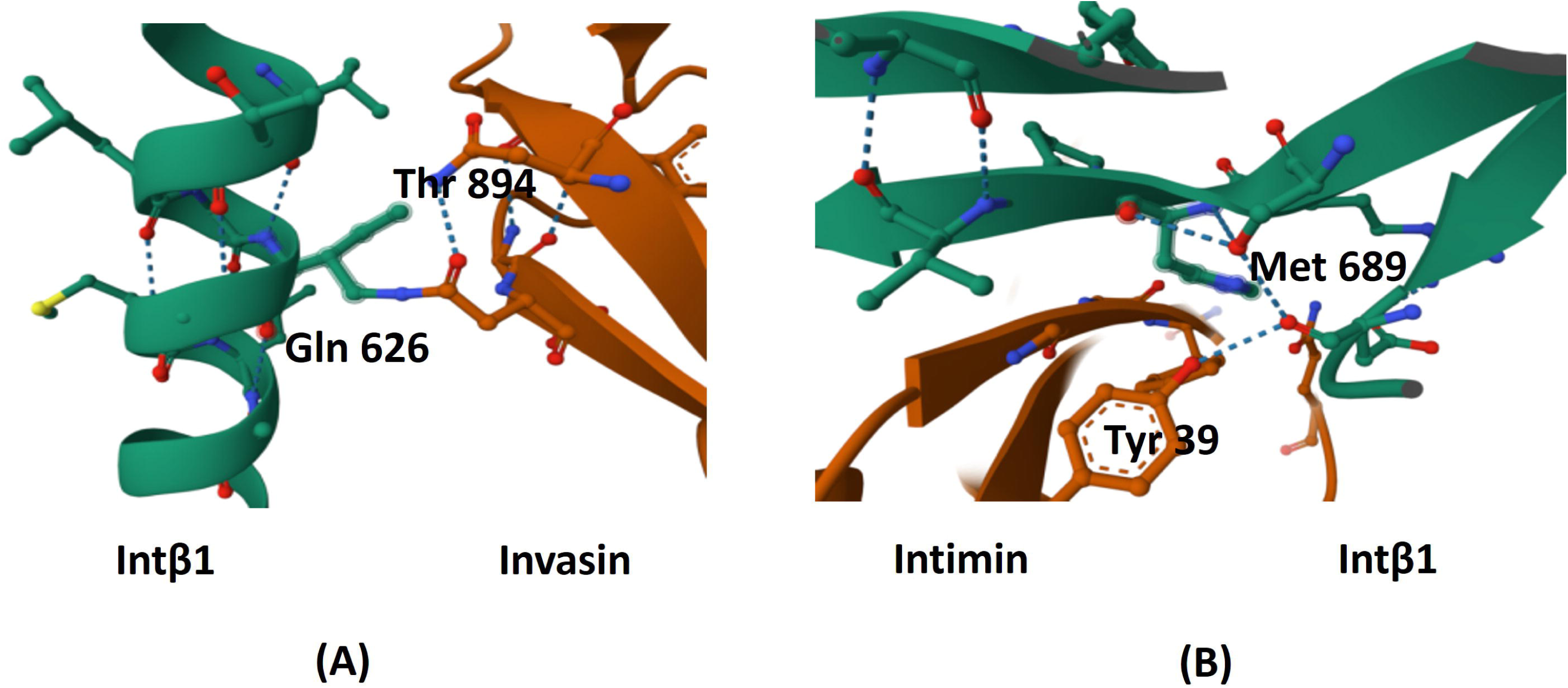
Example of interacting amino acid residues within the docking models (A) Thr 894 of invasin (D5) interacts with Gln 626 of integrin β14 and (B) Tyr 39 of intimin (D3) interacts with Met 689 of integrin β1.

## 4. Discussion

### 4.1. Invasin and intimin are not common in A. hydrophila

Invasins and intimins are proteins found in various bacterial species that cause attaching and effacing lesions on host cells [28]. These lesions result from the bacteria adhering to epithelial cells, disrupting the microvilli structure, and leading to changes in the cell surface that facilitate bacterial attachment and invasion. These species include *Hafnia alvei*, *Citrobacter rodentium*, and enteropathogenic and enterohaemorrhagic *Escherichia coli*, and enteropathogenic *Yersiniae* [29]. In silico study by previous workers showed that invasin and intimin proteins are predominantly associated with organisms from the γ-proteobacteria [30]. However, the tBLASTn results (Table S2) and synteny analysis (Fig. 1) clearly demonstrates that autotransporter type-3 (invasin and intimin) are only present in six strains (ATCC 7966, GSH8-2, LP0103, WCX23, WP8-S18-ESBL-02, and 23-C-23) out of 53 strains available for this study. This limited distribution suggests that these proteins may not be universally conserved across all *A. hydrophila* strains, indicating potential variability in pathogenicity or functional roles among different isolates. Interestingly, all thee strains having invasin and intimin were reported as hypervirulent strains [31].

### 4.2. Invasin and intimin are encoded in a single operon of A. hydrophila

Based on sequence differences, several distinct intimin orthologs have been identified. Tsai et al. (2010) reported that *A. hydrophila* carries an Ahy1 protein, which is classified within the invasin/intimin family, playing a role in bacterial adherence and invasion of host cells [30]. In this study, we discovered that the invasin and intimin proteins in *A. hydrophila* are encoded in a single operon on the bacterial chromosome. Notably, intimin is encoded by the yeeJ gene (Fig. 1). YeeJ was previously identified in *E. coli* as an inverse autotransporter that binds to peptidoglycan in the bacterial cell wall, a function crucial for biofilm formation, which allows bacteria to adhere to surfaces and resist environmental stressors. YeeJ has been shown to play a significant role in bacterial persistence, particularly in biofilm development [32], which is critical for pathogenicity and environmental survival. In *E. coli*, around 35% of the reported genomes carry the YeeJ gene [33], indicating its widespread presence and importance in bacterial ecology and infection mechanisms.

### 4.3. Invasin and intimin of A. hydrophila and β-integrins of Rohu are physiologically stable

The proteins examined in this study were largely uncharacterized, prompting an investigation into the important physicochemical properties. *A. hydrophila* invasin is composed of 916 amino acids with a molecular weight (MW) of 99,614.99 Da, while intimin comprises 535 amino acids with an MW of 56,335.50 Da. For comparison, Yersinia pseudotuberculosis produces an invasin that is a 986-amino-acid protein encoded by the chromosomal inv locus [34]. Whereas *In E. coli* K12 strains, full-length variants of intimin are expressed as intimin-α (939 amino acids), intimin-β (936 amino acids), or intimin-γ (934 amino acids) [35]. The physicochemical properties indicate stability and structural integrity. For example, *A. hydrophila* invasin exhibits a high aliphatic index (AI) of 83.08 and intimin shows an AI of 85.83, indicating a high degree of hydrophobicity, which contributes to protein stability. In contrast, both proteins display low instability indices (II) (Intimin: 25.02; Invasin: 29.18), which suggest that they are stable proteins. Additionally, both proteins have a negative GRAVY score, indicating they are hydrophilic overall, and exhibit acidic pI values (Intimin: 4.42; Invasin: 4.77), reinforcing their stability (Table 1).

The molecular cloning and sequence characterization of β1 integrin from *Cyprinus carpio* have been previously reported [36], yet there is limited information available regarding the molecular structure of fish integrins [37]. This study provides a comprehensive physicochemical characterization of all the β-integrins in Rohu. The β-integrins in Rohu exhibit comparatively low AI values (β2: 70.90 to β6: 78.31) and high instability indices (β2: 32.88 to β5b: 46.39). Similar to the test proteins, they also show negative GRAVY scores but have higher pI values ranging from 5.25 (β3) to 6.56 (β1) (Table 2). A protein with an instability index smaller than 40 is generally predicted to be stable, indicating that while some β-integrins may be less stable, they still exhibit features that could suggest functionality within their physiological context.

### 4.4. Invasin and intimin of A. hydrophila and β-integrins of Rohu are structurally stable

Intimin is characterized by the absence of disulfide bonds, while invasin contains a single disulfide bond, as outlined in Tables S4 and S5. In this study, the *A. hydrophila* invasin and intimin were found to exhibit a composition of α-helices, extended strands, and random coils, yet notably lacked any β-turns (Fig. 2A). This observation aligns with previous findings on the crystal structure of intimin from enteropathogenic *E. coli*, which similarly demonstrated a predominance of α-helices and extended strands without β-turns [38]. The 3D structures of invasin and intimin were modelled using templates A0A3T1A166 and A0A4P7IWM0, derived from *A. hydrophila* strains ATCC 7966 and WCX23 (Table S7). These models are likely the closest approximations available, supported by validation from the ProSA and ProQ servers. Experimental evidence from invasion of *Yersiniae* and *Escherichia coli*, shows a topology comprising a 12-stranded β-barrel with an α-helical linker located in the pore of the barrel, in analogy to classical Ats but with an inverted N-to C-terminal domain order [39]. The crystal structures of enterohemorrhagic *E. coli* and *Y. pseudotuberculosis*, the translocation units of both intimin and invasion [17] confirmed this topology model of a 12-stranded β-barrel with an α-helical linker residing in the largely hydrophilic pore.

In eukaryotes, integrins represent a diverse family of cell adhesion receptors that play a crucial role in various biological processes, including cell signalling, immune responses, and tissue repair. This family comprises eight β-subunits and eight α-subunits, which non-covalently associate to form 24 distinct αβ integrin heterodimers. Each β subunit can bind to multiple α subunits, allowing for a vast array of integrin combinations. Both the α-and β-subunits are type I transmembrane receptors that have a cytoplasmic tail, a single transmembrane domain, and a sizable extracellular region in common [40]. All β integrins contain an inserted domain (I domain), which is homologous to the von Willebrand factor A domain (vWFA). In the phylogenetic tree based on sequences of the β integrins, the vertebrate sequences constitute two major branches. Group A contains β1, β2, and β7 and group B contains β3, β5, β6, and β8 [41]. Fish integrins are relatively less studied compared to mammalian integrins, particularly in humans. No crystal structures are available for Rohu β-integrins. Huhtala et al. (2005) identified eight β integrins in pufferfish, noting the absence of β2 and β7 orthologues of human β integrins, while observing duplications of β1 and β3 orthologues [42]. In contrast, our current findings reveal that the eight β integrins of Rohu are indeed orthologs of human β integrins, suggesting a more complete representation of integrin diversity in this species.

### 4.5. Docking interactions between invasin/intimin of A. hydrophila and β-integrins of Rohu

High-affinity binding of invasin is necessary for bacterial internalization [43]. Invasin may recognize an un-glycosylated region of integrins [44]. According to Leong et al., (1993), the disulfide bond between cysteines is required for integrin binding as because it is necessary for proper folding [45]. Hamburger et al. (1999) showed that aspertate residue in invasin D5 in *Y. psudotuyberculosis* are required for integrin binding [44]. In present investigation, *A. hydrophila* ATCC 7966 invasin D5 contains 5 aspertate residues, D4 consists of 8, while D3 contains highest numbers of aspertate residues (17 residues), D2 contains 9 and 6 residues are present in the D1 (Table 1). The overall similarity in the relative positions of these residues suggests that invasin and host proteins share common integrin-binding features. On the same way, *A. hydrophila* ATCC 7966 intimin D3 contains 8 aspertate residues, D2 contains 10, and D1 contains 7, and 6 aspertate residues are present in D0.

A more negative docking score indicates a stronger potential binding affinity between the two molecules. Generally, a docking score around –200 or lower is considered favorable for interaction. In the current study, the docking scores for the experimental models ranged from –405.89 to –246.23, reflecting strong binding potential. Additionally, the confidence scores, which ranged from 0.8734 to 0.9832, further support the likelihood of binding; scores above 0.7 suggest a high probability of interaction. These findings indicate that both invasin and intimin from *A. hydrophila* exhibit a robust binding affinity to all β-integrin members of Rohu fish. Such high-affinity interactions are likely crucial for the bacteria’s attachment to the extracellular matrix and for facilitating cell–cell interactions with the host [46]. Previous studies have also demonstrated similar tight binding, such as the interaction of invasin and intimin from *Yersinia enterocolitica* with β1 integrin in humans, underscoring the importance of these adhesin proteins in bacterial pathogenesis [47]. This highlights the evolutionary conservation of binding mechanisms among different pathogens targeting integrin receptors, potentially aiding in the development of therapeutic strategies against bacterial infections in both aquatic and terrestrial hosts.

## 5. Conclusion

The limited distribution of invasin and intimin among hypervirulent strains of *A. hydrophila*, combined with their strong binding affinity to all eight β-integrins of Rohu, underscores the significant role these type-3 autotransporters in disease establishment. This specificity suggests that only certain strains of *A. hydrophila* possess the necessary tools to effectively interact with host cells, facilitating adherence and invasion.

## CRediT authorship contribution statement

**Agradip Bhattacharyya:** Data curation, Formal analysis, Methodology, Writing original draft, Software, Visualization. **Goutam Banerjee:** Conceptualization, Software, Formal analysis, Writing original draft, Review and editing, Visualization. **Pritam Chattopadhyay:** Conceptualization, Software, Data acquision, Formal analysis, Methodology, Writing original draft, Review and editing, Visualization, Supervision

## Declaration of competing interest

None of the authors have any conflict of interest.

## Data availability

All data are available in public databases like KEGG Orthology database (https://www.genome.jp/kegg/ko.html), UniProt database (https://www.uniprot.org/), and NCBI Genome database (https://www.ncbi.nlm.nih.gov/genome)

## Acknowledgements

We are grateful to Raja Rammohun Roy Mahavidyalaya and M.U.C. Women’s College, India, as well as the University of Illinois, Urbana-Champaign, Illinois, USA, for providing the necessary resources and support for this study. This research received no external funding.

## Notes

https://www.genome.jp/kegg/ko.html

https://www.uniprot.org/

https://www.ncbi.nlm.nih.gov/genome

### Competing Interest Statement

The authors have declared no competing interest.

