## Supplementary for "Investigating the potential role of intimin/invasin in Aeromonas hydrophila virulence in Labeo rohita: a host-microbe interaction study"

**Table S1:**The accession numbers and DNA and protein sequences of the test proteins (Invasin, and Intimin) of *A. hydrophila* ATCC 7966

| Test Proteins | NCBI Protein ID | UniProt ID | DNA sequence | Protein sequence |
|---|---|---|---|---|
| **Invasin** | ABK38065 | A0KH56 | atgaacttcacaccaagcaaagtccatcagttttttgga<br>cttatctctttttaggacagcaggcattatttcctgtggtgt<br>atgcggctggatcactctttacattcgcttctgagcaacc<br>tgaacttatccgctatctcatcaaaaagggagatacgctt<br>gattctgtctcacgccagtttggggtggtgccacatcaa<br>ctgttgacattaaaccgccacctgaaactcgatgcacct<br>tatctggtggttggggagaccataaatgtaccacgacc<br>acagaccttgcccatattagttagcgagaatccactgta<br>cctgaaagaaaatacagagagtcacattaatgtggaac<br>aaaaagtggctcagcatgttactcgcctggggcaggca<br>ctcacagcacttgaagaggggagggaggctggttttga<br>aaatgagagtgagatagttaaaggtgcgcgtcgtgcac<br>agcgcaatagcatagctgcctctggagagcaggtgcc<br>gacatctgcatcgcgctatggctcggagcaggaggtg<br>cagtactggcgtcagcagctcgcgacacagtttgaaga<br>ggaggccaatgcgtatgctgcatccttgctgggggcga<br>tgggcacagccagaacccgggtgacgctggatgacg<br>atttcaatatggtcactgccgaggcggatctcttgttgcc<br>gcttgctgaagagcagcagactctgttgtttacccagttt<br>ggtctacgccgaaatgggcaggatcgtaccattgccaa<br>tctgggggtgggccagcgtcatttcctcgaccgctgga<br>tgctgggttacaacctgtttgctgattatgacctgaccaat<br>cggcactggcgtgcgggagtcggagctgaggcgtgg<br>cgtgattatctcaagttgggcgccaacttttataccccac<br>tgagcagttggcgtgactccccacggtttgagggtatg<br>gaagagcgcgcagcacgcggcatggatgtgcgtcttg<br>aggcctatctgccagcctatccgcaatggagcgcatcc<br>ctcactgctgagcagtatttgggggagcgggttggcttg<br>ctggacgctgatcagcttgagcgcgatccgcatgccat<br>tactgctggtctgcactataaccccttcccgttactcaag<br>atggatgtcgagcaagttgaagcgagcggcaggcaac<br>atgacacccgctttacgctggggctggagtggaagctc<br>ggggcaacgctctgggacatgctgaacccgtccagtg<br>tcgacaagtcgttggcgggtatgcgccacgatctgattg<br>agcgtaacaatgacatggtgcttgagtatcgtgacaag<br>gtgctcctcaaggcgtctctcaacgaccaatattccgcg<br>gttgaggggcaagcactgaccttgacgctcaacatcca<br>gcattcccgtcagatagcctccatccagtggttgggtga<br>cgtcctggggctcagtgggcttagcccggctgatacgg<br>ctgggcaggacaaacgcgccctcacactcccgtccct<br>gcccacctatcgtattggccaatccaatcaatatccggt<br>cgttgccatcgtcaccgacatcgatggccatgaagcca<br>ttgcggaaggggttgttgccgtttcggaagactctgggc<br>tgcaacctgctatccagttggcagagcactttgttcaact<br>tctacccggtgctcactaccaagttgactgggggttgt<br>tgacccccgtcagaagcccgccaaagcacgcaatgg<br>gatcccgattgagagcattgagggggatgttgaagcg | MNFTPSKVHQFFWT<br>YLFLGQQALFPVVYA<br>AGSLFTFASEQPELIR<br>YLIKKGDTLDSVSRQ<br>FGVVPHQLLTLNRHL<br>KLDAPYLVVGETINV<br>PRPQTLPILVSENPLY<br>LKENTESHINVEQKV<br>AQHVTRLGQALTALE<br>EGREAGFENESEIVK<br>GARRAQRNSIAASGE<br>QVPTSASRYGSEQE<br>VQYWRQQLATQFEE<br>EANAYAASLLGAMG<br>TARTRVTLDDDFNM<br>VTAEADLLLPLAEEQ<br>QTLLFTQFGLRRNGQ<br>DRTIANLGVGQRHFL<br>DRWMLGYNLFADYD<br>LTNRHWRAGVGAEA<br>WRDYLKLGANFYTP<br>LSSWRDSPRFEGMEE<br>RAARGMDVRLEAYL<br>PAYPQWSASLTAEQY<br>LGERVGLLDADQLE<br>RDPHAITAGLHYNPF<br>PLLKMDVEQVEASG<br>RQHDTRFTLGLEWK<br>LGATLWDMLNPSSV<br>DKSLAGMRHDLIER<br>NNDMVLEYRDKVLL<br>KASLNDQYSAVEGQ<br>ALTLTLNIQHSRQIAS<br>IQWLGDVLGLSGLSP<br>ADTAGQDKRALTLPS<br>LPTYRIGQSNQYPVV<br>AIVTDIDGHEAIAEG<br>VVAVSEDSGLQPAIQ<br>LAEHFVQLLPGAHY<br>QVDWGVVDPRQKPA<br>KARNGIPIESIEGDV<br>EADRDGYQYRYRLT<br>FQGPGQVLTGRELVA<br>PGQPNPGTRYRLDVE<br>VIFPSGHVARDSMEF<br>EFIDDATVPGAPTLTA<br>ADSNGDDKPEVTGK |

| | | | | |
|---|---|---|---|---|
| | | | gaccgcgatggttaccagtaccgctatcgcctgacatttcaagggcctggacaggtactgacggggcgtgagctggttgcacccgggcaacccaaccccggcacgcggtatcgcttggatgttgaggtcatcttcccgagtggtcatgtggcgcgtgacagcatggagtttgagtttattgatgacgcgacggtgccgggggccccgacactgacagcggccgacagcaatggtgacgacaagccggaggtcactggtaaagcggaaccagagagcacggtgaccattacctggccggatggctcgaccagcaccacgactgcagatgtggatggcaactacaccctggaagctccgacggtgcaggggagtggcaccatcaccgccacggctaccgacaagagcggcaacaccgggccggctaccagcgtgaactacatcgactcaacggtaccgggggccccgacgttggctgcaacggatagcaatagcgacaacaagccggaggtcagtggcaaggcagagccggacagcacggtgaccattacctggccggatggcacaaccagcaccacggctgcagatgtggatggcaactacaccctggaagctccgacggtgcaggggagtggcaccatcaccgccacggctaccgacaagagcggcaacactgggcctgctagcagtgtgaactatctcgcgagcttggtgggggtgacgattaccgggctggttgacggctttcctcaggtgggagcggtactgacggccgtgtctgactgcggctcgagcgcctgccgtgcaggactgacctggcagtggcagattgaagatggtgtcggcagtggaaattttgtggatatcggtggggcgaccagcgtgacctatttacccgcgcgagaagatcagaagaaacgggtgcgcgtcattgttgcgccgctgtga | AEPESTVTITWPDGSTSTTTADVDGNYTLEAPTVQGSGTITATATDKSGNTGPATSVNYIDSTVPGAPTLAATDSNSDNKPEVSGKAEPDSTVTITWPDGTTSTTAADVDGNYTLEAPTVQGSGTITATATDKSGNTGPASSVNYLASLVGVTITGLVDGFPQVGAVLTAVSDCGSSACRAGLTWQWQIEDGVGSGNFVDIGGATSVTYLPAREDQKKRVRVIVAPL |
| **Intimin** | **ABK37862** | **A0KH58** | atggccgaaagtgccaacaccctgtcggtgcaggggcgtgcgccagtggcatctggcgtgacactgactggcccggcggcgccattggtgggggataccctgctgggaacatatacctttaatgatctggatgccgatgaggaagagggcagtgtattgcgttggctaaataatagtgacatcgaactcgccagaggtgatcgctatctgttgaccgatgccgaacgtggcaaacagcttcgctttgctgttaccccggccaccaatccggcggtgactgatccgcatgagggggcagaggtaagcagttctctctctgcggtggtgcaggggcggccagaccccgccaaatccaccttcagtgccaacaagtccaccatagtggcggatggggttgataacgcagttctgacgctgaccctcaaagacgacaatcaagcaccggtaaccggcatcagtgatcgggttgcgctaggctacagcggtgtggatacagatgttatctccctgactgaaaacgatctgggcaatggcatctatgagtacatcctgacggggaccaagtctggcgttgtcactctgacgccacagctggatggcgctcccttgaccactatccctcaatcactgcaggtgactctggccgctaacccagccaccacgaaagtggtccaactggtgaccacaacagacagtcagcctgcggataacgtcagcgctgacctgctgacagcgactgtgcatgatattgaggggaaccccatgcagggcgctagcgtggtgtggtcgcatgggagcaccacggcaaccttgggcgcagcgacaagcgtgaccgacagtaatggtctggcaacggtgagcctcatcaacacccgggctgagacagtcgccgtgacggccagggttcaagccaattccggtgactcgggcatgaccagtgatgccaacttcgcactctatccggtgctcgatacattagccgtagggaccaataatcagcccgccaataataaggccttgaacacaatagtggccacattcaaagaccttgatggggcggttctggcaaatgtgccgatcactgtgcagttcactgctga | MAESANTLSVQGRAPVASGVTLTGPAAPLVGDTLLGTYTFNDLDADEEEGSVLRWLNNSDIELARGDRYLLTDAERGKQLRFAVTPATNPAVTDPHEGAEVSSSLSAVVQGRPDPAKSTFSANKSTIVADGVDNAVLTLTLKDDNQAPVTGISDRVALGYSGVDTDVISLTENDLGNGIYEYILTGTKSGVVTLTPQLDGAPLTTIPQSLQVTLAANPATTKVVQLVTTTDSQPADNVSADLLTATVHDIEGNPMQGASVVWSHGSTTATLGAATSVTDSNGLATVSLINTRAETVAVTARVQANSGDSGMTSDANFALYPVLDTLAVGTNNQPANNKALNTIVATFKDLDGAVLANVPITVQFTADKSTVTFDNESPWTTTTNDAGAVIVSLRNNVVENVEVTLYATNSSTVQKVASVNFTELMISRFLKP |

| | | | | |
|---|---|---|---|---|
| | | | caagagcactgtcacgttcgataacgagtcgccatgga cgaccactaccaatgacgctggtgcggtgatcgtgtca ctgcgtaacaacgtggttgagaatgttgaggttacgcta tatgcgacaaacagcagtacagtgcaaaaagtggcctc ggttaactttacagaattgatgatatcgagattcctcaag ccggatccggccaccatgaattggctagatgccgaca actattgcaataacatcgggcgccgtttaccgacggtg gatgaattgagagcattgtttgtcgaggtgtccccttcctt tggtcagaattatgaattgtgttatgttcatggatggccaa tggatggcaagtgtggtggtacctacgatgactattgga cgagtgagaagtatgttccccatggtactagcgctcatg ccgccgtctatatgcacaccggtgctgttggtggattcg gtgacacacagcctagtcaggtagtatgcatacgtcgtt ga | DPATMNWLDADNYC NNIGRRLPTVDELRA LFVEVSPSFGQNYEL CYVHGWPMDGKCG GTYDDYWTSEKYVP HGTSAHAAVYMHTG AVGGFGDTQPSQVV CIRR |

**Table S2:** Results of microbial genome BLAST for Invasin and Intimin within the *A. hydrophila* genomes using the tBLASTn algorithm

| Sl. No. | *A. hydrophila* strains | Invasin | Intimin |
|---|---|---|---|
| 1 | 2359 | 0 | 0 |
| 2 | 23-C-23 | 1729 | 953 |
| 3 | 3019 | 0 | 0 |
| 4 | 3206 | 0 | 0 |
| 5 | 3924 | 0 | 0 |
| 6 | 4960 | 0 | 0 |
| 7 | 4AK4 | 0 | 0 |
| 8 | 71317 | 0 | 0 |
| 9 | 71339 | 0 | 0 |
| 10 | A008N2 | 0 | 0 |
| 11 | AC133 | 0 | 0 |
| 12 | AC185 | 0 | 0 |
| 13 | Aer_Brac14A | 0 | 0 |
| 14 | Aer_Brac66 | 0 | 0 |
| 15 | Aer_Pi25.1HTAS | 0 | 0 |
| 16 | AH10 | 0 | 0 |
| 17 | Ah2111 | 0 | 0 |
| 18 | Ah27 | 0 | 0 |
| 19 | AHNIH1 | 0 | 0 |
| 20 | AL06-06 | 0 | 0 |
| 21 | AL09-71 | 0 | 0 |
| 22 | ATCC 7966 | 1846 | 1049 |
| 23 | B11 | 0 | 0 |
| 24 | Brac6 | 0 | 0 |
| 25 | CSUSB2 | 0 | 0 |
| 26 | D4 | 0 | 0 |
| 27 | FDAARGOS_916 | 0 | 0 |
| 28 | GSH8-2 | 1846 | 1046 |
| 29 | GYK1 | 0 | 0 |
| 30 | HX-3 | 0 | 0 |
| 31 | J-1 | 0 | 0 |
| 32 | JBN2301 | 0 | 0 |
| 33 | K522 | 0 | 0 |
| 34 | KAM330 | 0 | 0 |
| 35 | KN-Mc-1R2 | 0 | 0 |

| | | | |
|---|---|---|---|
| 36 | LHW39 | 0 | 0 |
| 37 | LP0103 | 1837 | 1006 |
| 38 | ML09-119 | 0 | 0 |
| 39 | MX16A | 0 | 0 |
| 40 | NEB724 | 0 | 0 |
| 41 | NJ-35 | 0 | 0 |
| 42 | NN-MR659 | 0 | 0 |
| 43 | NUITM-VA1 | 0 | 0 |
| 44 | OnP3.1 | 0 | 0 |
| 45 | PartN-Ahydrophila-RM8376 | 0 | 0 |
| 46 | pc104A | 0 | 0 |
| 47 | WCHAH045096 | 0 | 0 |
| 48 | WCX23 | 1729 | 953 |
| 49 | WP7-S18-ESBL-06 | 0 | |
| 50 | WP8-S18-ESBL-02 | 1846 | 1046 |
| 51 | YL17 | 0 | 0 |
| 52 | ZYAH72 | 0 | 0 |
| 53 | ZYAH75 | 0 | 0 |

**Table S3:** Secondary structure prediction of Invasin and Intimin of *A. hydrophila* ATCC 7966

| Composition | Alpha-helix | Extended strand | β -turn | Random coil |
|---|---|---|---|---|
| **Invasin** | 9.72% | 25.66% | 0.00% | 64.63% |
| **Intimin** | 0.19% | 29.35% | 0.00% | 70.47% |

**Table S4:** Secondary structure prediction of β-integrins of Rohu (*Labeo rohita*)

| Composition | Alpha-helix | Extended strand | β -turn | Random coil |
|---|---|---|---|---|
| β1 | 20.47 % | 14.73% | 0.00% | 64.79% |
| β2 | 20.99% | 15.12% | 0.00% | 63.89% |
| β3 | 21.46% | 15.96% | 0.00% | 62.58% |
| β4 | 11.40% | 23.99% | 0.00% | 64.61% |
| β5 | 20.37% | 16.27% | 0.00% | 63.35% |
| β6 | 21.69% | 17.46% | 0.00% | 60.85% |
| β7 | 22.27% | 16.02% | 0.00% | 61.72% |
| β 8 | 23.16% | 14.46% | 0.00% | 62.38% |

**Table S5**: Functional Analysis Invasin and Intimin of *A. hydrophila* ATCC 7966

| Test Proteins | No of Cysteine Amino acids | No. of Di-sulphide bonds |
|---|---|---|
| **Invasin** | 2 | 1 |
| **Intimin** | 4 | 0 |

**Table S6:** Functional Analysis of β-integrins of *Rohu (Labeo rohita)*

| β-integrins | No of Cysteine Amino acids | No. of Di-sulphide bonds |
|---|---|---|
| β1 | 58 | 24 |
| β2 | 59 | 22 |
| β3 | 56 | 32 |
| β4 | 64 | 22 |
| β5 | 59 | 26 |
| β6 | 57 | 25 |
| β7 | 55 | 26 |
| β 8 | 55 | 22 |

**Table S7:** Homology modeling data of Invasin and Intimin of *A. hydrophila* by SWISS-MODEL accessed via the ExPASy web server

| Component | Template | Description | Oligostate | Sequence similarity | GMQE |
|---|---|---|---|---|---|
| **Invasin** | A0A3T1A166.1.A | UniProtKB entry unknown, most likely obsolete AlphaFold DB model of A0A3T1A166 (gene: unknown, organism: unknown) | Monomer | 100% | 0.72 |
| **Intimin** | A0A4P7IWM0.1.A | UniProtKB entry unknown, most likely obsolete AlphaFold DB model of A0A4P7IWM0 (gene: unknown, organism: unknown) | Monomer | 89.91% | 0.91 |

**Table S8:** Homology modeling of the β-integrins of Rohu (*Labeo rohita*) by SWISS-MODEL accessed via the ExPASy web server

| Integrins | Model | Description | Oligostate | Sequence similarity | GMQE |
|---|---|---|---|---|---|
| β1 | A0A286Y9W0.1.A | Integrin beta AlphaFold DB model of A0A8M6YW86_DANRE (gene: itgb1a, organism: Danio rerio (Zebrafish) (*Brachydanio rerio*)) | Monomer | 95.01% | 0.87 |
| β2 | E7FCN5.1.A | Integrin beta AlphaFold DB model of E7FCN5_DANRE (gene: itgb2, organism: Danio rerio (Zebrafish) (Brachydanio rerio) | Monomer | 80.57% | 0.86 |

| | | | | | |
|---|---|---|---|---|---|
| β3 | B3DIP9.1.A | UniProtKB entry unknown, most likely obsolete<br>AlphaFold DB model of B3DIP9 (gene: unknown, organism: unknown) | Monomer | 95.06% | 0.84 |
| β4 | F1RA51.1.A | UniProtKB entry unknown, most likely obsolete<br>AlphaFold DB model of F1RA51 (gene: unknown, organism: unknown) | Monomer | 86.55% | 0.71 |
| β5 | F1QF91.1.A | UniProtKB entry unknown, most likely obsolete<br>AlphaFold DB model of F1QF91 (gene: unknown, organism: unknown) | Monomer | 95.26% | 0.83 |
| β6 | F1QGX0.1.A | Integrin beta<br>AlphaFold DB model of F1QGX0_DANRE (gene: itgb6, organism: Danio rerio (Zebrafish) (*Brachydanio rerio*) | Monomer | 88.83% | 0.84 |
| β7 | E7F4H9.1.A | UniProtKB entry unknown, most likely obsolete<br>AlphaFold DB model of E7F4H9 (gene: unknown, organism: unknown) | Monomer | 75.13% | 0.84 |
| β 8 | A0A7J6C0U9.1.A | Integrin beta<br>AlphaFold DB model of A0A7J6C0U9_9TELE (gene: A0A7J6C0U9_9TELE, organism: *Onychostomamacrolepis*) | Monomer | 89.61% | 0.73 |

**Table S9:** Evaluation of tertiary structure of Invasin, Intimin and BamAof *A. hydrophila*by SWISS-MODEL accessed via the ExPASy web server

| | **Ramachandran plot analysis** | | **ProQ** | | **ProSA** |
|---|---|---|---|---|---|
| **Protein** | **Residues in the favored region (%)** | **Residues in the Outlier region(%)** | **LG score** | **Maxsub** | **Z score** |
| **Invasin** | 94.31% | 0.88 % | 11.489 | -0.871 | -7.21 |
| **Intimin** | 95.16 % | 0.27 % | 10.667 | -1.691 | -8.67 |

**Table S10:** Evaluation of tertiary structure of β-integrins of Rohu (*Labeo rohita*) by SWISS-MODEL accessed via the ExPASy web server

| | Ramachandran plot analysis | | ProQ | | ProSA |
|---|---|---|---|---|---|
| **Protein** | **Residues in the favored region (%)** | **Residues in the Outlier region(%)** | **LG score** | **Maxsub** | **Z score** |
| β1 | 96.25 | 0.75 | 9.647 | -0.548 | -10.55 |
| β2 | 96.60 | 0.65 | 11.038 | -0.659 | -10.65 |
| β3 | 96.07 | 0.63 | 11.218 | -0.751 | -09.7 |
| β4 | 87.71 | 4.88 | -0.835 | -0.113 | -13.47 |
| β5 | 94.27 | 1.62 | 10.442 | -0.594 | -10.78 |
| β6 | 95.11 | 1.54 | 9.884 | -0.517 | -10.54 |
| β7 | 94.46 | 0.92 | 10.113 | -0.599 | -10.64 |
| β 8 | 92.80 | 1.60 | 9.409 | -0.463 | -10.6 |

**Table S11:** Evaluation of receptor-ligand interface residue pair(s)

| **Receptors (Rohu Integrins)** | **Invasin** | | **Intimin** | |
|---|---|---|---|---|
| | **Interacting Amino Acids** | **RMSD*** | **Interacting Amino Acids** | **RMSD*** |
| **Integrin β1** | 564A - 842A | 4.227 | 6A - 177A | 4.413 |
| | 598A - 869A | 2.474 | 13A - 142A | 3.606 |
| | 599A - 869A | 4.475 | 17A - 117A | 4.751 |
| | 600A - 869A | 3.195 | 17A - 142A | 2.659 |
| | 601A - 869A | 4.494 | 17A - 146A | 3.971 |
| | 617A - 844A | 4.887 | 20A - 116A | 4.514 |
| | 617A - 858A | 3.522 | 564A - 30A | 2.836 |
| | 617A - 860A | 4.601 | 564A - 31A | 2.946 |
| | 626A - 894A | 4.186 | 564A - 112A | 3.853 |
| | 626A - 895A | 3.079 | 565A - 73A | 2.910 |
| | 626A - 896A | 4.469 | 565A - 76A | 2.631 |
| | 627A - 896A | 3.758 | 565A - 144A | 2.188 |
| | 628A - 869A | 3.445 | 566A - 112A | 2.991 |
| | 628A - 896A | 4.977 | 566A - 113A | 4.049 |
| | 629A - 869A | 2.283 | 566A - 115A | 4.435 |
| | 631A - 869A | 4.576 | 566A - 203A | 4.607 |
| | 676A - 873A | 2.569 | 571A - 201A | 4.487 |
| | 676A - 914A | 3.423 | 617A - 68A | 3.011 |
| | 676A - 915A | 2.381 | 618A - 68A | 2.952 |
| | 676A - 916A | 2.841 | 620A - 68A | 4.074 |
| | 677A - 835A | 2.865 | 625A - 67A | 4.838 |
| | 677A - 915A | 3.149 | 626A - 35A | 4.395 |
| | 677A - 916A | 3.900 | 626A - 36A | 2.820 |
| | 679A - 916A | 4.023 | 626A - 37A | 3.148 |
| | 681A - 812A | 4.481 | 626A - 38A | 4.461 |
| | 681A - 835A | 4.033 | 626A - 53A | 4.807 |
| | 681A - 870A | 3.293 | 626A - 67A | 3.760 |
| | 681A - 871A | 4.612 | 626A - 68A | 4.900 |
| | 681A - 916A | 3.800 | 627A - 35A | 3.253 |

| | | |
|---|---|---|
| | 682A - 870A 3.237<br>682A - 871A 4.578<br>682A - 916A 4.428<br>683A - 870A 3.495<br>690A - 916A 3.322<br>735A - 259A 4.320<br>736A - 287A 4.289<br>738A - 259A 3.681<br>739A - 259A 3.641<br>739A - 278A 2.068<br>739A - 279A 4.856<br>739A - 280A 3.835<br>739A - 287A 2.206<br>740A - 287A 3.041<br>741A - 12A 2.477<br>742A - 12A 3.869<br>742A - 15A 2.352<br>742A - 245A 2.639<br>742A - 257A 3.398<br>742A - 259A 3.881<br>742A - 282A 4.284<br>743A - 257A 4.337<br>743A - 280A 3.809<br>743A - 282A 1.916<br>744A - 282A 3.427<br>745A - 12A 4.252<br>745A - 16A 2.745<br>746A - 245A 4.635<br>746A - 247A 1.733<br>746A - 255A 4.253<br>746A - 256A 4.362<br>746A - 257A 3.248<br>746A - 282A 2.701<br>747A - 282A 3.073<br>747A - 283A 4.581<br>749A - 16A 4.558<br>749A - 20A 4.315<br>749A - 24A 3.685<br>750A - 249A 3.901<br>752A - 20A 4.494<br>752A - 100A 4.156<br>753A - 24A 3.694<br>756A - 98A 4.215<br>756A - 99A 4.013<br>756A - 100A 3.352<br>756A - 101A 4.837<br>759A - 100A 3.448<br>760A - 28A 4.844<br>760A - 97A 2.566<br>760A - 98A 3.718<br>760A - 99A 2.520 | 627A - 68A 2.804<br>628A - 35A 4.795<br>628A - 68A 4.579<br>630A - 68A 4.957<br>634A - 37A 4.602<br>634A - 38A 1.807<br>634A - 39A 2.788<br>634A - 40A 2.922<br>635A - 38A 3.835<br>635A - 39A 4.795<br>635A - 40A 2.861<br>635A - 48A 1.811<br>636A - 39A 4.566<br>636A - 40A 3.267<br>636A - 41A 4.858<br>637A - 40A 3.319<br>637A - 41A 4.065<br>637A - 42A 3.741<br>637A - 45A 4.956<br>637A - 46A 3.482<br>637A - 47A 4.670<br>637A - 48A 4.831<br>638A - 40A 3.797<br>638A - 41A 3.111<br>638A - 42A 3.241<br>638A - 43A 4.297<br>639A - 41A 2.879<br>640A - 18A 2.557<br>640A - 39A 3.991<br>640A - 41A 2.465<br>641A - 16A 3.003<br>641A - 41A 2.763<br>641A - 42A 4.711<br>641A - 43A 3.629<br>642A - 43A 3.213<br>676A - 21A 3.912<br>676A - 104A 3.616<br>689A - 39A 2.430<br>690A - 18A 4.108<br>690A - 19A 3.558<br>690A - 39A 3.262<br>691A - 18A 3.526<br>691A - 19A 3.511<br>692A - 17A 3.896<br>692A - 18A 3.209<br>692A - 19A 2.718<br>692A - 20A 3.912<br>692A - 101A 4.515<br>692A - 102A 3.213<br>692A - 103A 3.851<br>692A - 104A 3.731<br>693A - 16A 3.318<br>693A - 17A 4.643<br>693A - 18A 4.158<br>694A - 100A 2.523<br>694A - 101A 3.514<br>694A - 102A 4.533<br>700A - 101A 3.025<br>700A - 102A 3.865 |

| | | |
|---|---|---|
| **Integrin β2** | 1A - 12A 4.379<br>2A - 287A 2.933<br>3A - 15A 4.833<br>3A - 245A 4.256<br>3A - 257A 2.925<br>3A - 258A 4.353<br>3A - 259A 4.133<br>3A - 278A 4.050<br>3A - 280A 3.252<br>3A - 287A 3.198<br>4A - 259A 4.448<br>6A - 278A 4.052<br>6A - 287A 3.615<br>6A - 310A 3.584<br>7A - 259A 3.512<br>7A - 260A 4.055<br>7A - 261A 3.332<br>7A - 276A 2.962<br>7A - 277A 3.174<br>7A - 278A 3.091<br>10A - 276A 2.196<br>10A - 277A 4.037<br>10A - 289A 3.272<br>10A - 290A 3.898<br>10A - 291A 3.716<br>10A - 308A 4.936<br>10A - 310A 4.237<br>11A - 261A 4.358<br>11A - 276A 2.691<br>14A - 276A 3.806<br>14A - 291A 3.694<br>543A - 873A 4.843<br>543A - 916A 4.241<br>544A - 872A 4.667<br>544A - 873A 4.097<br>544A - 894A 4.822<br>544A - 916A 2.597<br>545A - 872A 3.755<br>545A - 873A 2.348<br>545A - 874A 1.965<br>545A - 875A 3.681<br>545A - 894A 2.039<br>545A - 895A 4.796<br>546A - 875A 3.320<br>546A - 889A 4.970<br>546A - 894A 4.282<br>547A - 875A 4.102<br>547A - 892A 3.882<br>547A - 894A 2.895<br>572A - 916A 4.692<br>597A - 870A 4.367<br>605A - 816A 4.828<br>605A - 829A 4.187<br>613A - 818A 3.773<br>613A - 826A 4.790<br>613A - 827A 3.989<br>613A - 828A 4.900<br>613A - 829A 4.115 | 17A - 500A 3.267<br>20A - 439A 2.826<br>20A - 500A 3.560<br>20A - 507A 3.149<br>20A - 509A 2.541<br>20A - 524A 3.615<br>21A - 505A 3.886<br>21A - 506A 3.776<br>21A - 507A 1.844<br>21A - 524A 4.851<br>22A - 507A 4.017<br>23A - 506A 4.702<br>23A - 507A 2.578<br>23A - 508A 4.153<br>23A - 522A 4.112<br>23A - 523A 1.807<br>23A - 524A 2.342<br>23A - 525A 2.913<br>41A - 506A 3.401<br>41A - 526A 3.972<br>42A - 525A 4.216<br>64A - 525A 4.182<br>65A - 525A 3.204<br>65A - 526A 4.526<br>66A - 525A 4.234<br>69A - 438A 4.947<br>75A - 436A 4.363<br>75A - 527A 3.466<br>80A - 525A 4.712<br>80A - 526A 2.524<br>80A - 527A 4.768<br>81A - 526A 4.289<br>82A - 492A 3.970<br>82A - 494A 3.571<br>82A - 522A 3.708<br>82A - 526A 2.971<br>82A - 527A 2.229<br>82A - 528A 4.023<br>83A - 513A 2.565<br>83A - 521A 4.461<br>83A - 522A 4.728<br>84A - 513A 4.605<br>85A - 513A 4.904<br>85A - 515A 4.560<br>85A - 516A 3.831<br>85A - 518A 3.212<br>86A - 472A 4.908<br>87A - 516A 3.203<br>90A - 471A 2.872<br>90A - 472A 3.214<br>92A - 469A 4.959<br>112A - 526A 4.077<br>114A - 503A 4.128<br>114A - 504A 3.532<br>115A - 503A 4.522<br>115A - 504A 4.783<br>115A - 521A 3.591<br>115A - 522A 4.677 |

| | | |
|---|---|---|
| | 614A - 827A 4.401<br>614A - 828A 2.604<br>614A - 829A 2.733<br>616A - 753A 4.379<br>661A - 747A 2.492<br>661A - 748A 4.992<br>661A - 749A 2.672<br>669A - 747A 4.987<br>669A - 749A 3.389<br>701A - 742A 4.291<br>702A - 670A 4.656<br>702A - 742A 2.889<br>705A - 667A 4.842<br>705A - 740A 4.920<br>706A - 667A 3.814<br>709A - 667A 3.672<br>709A - 739A 4.881<br>713A - 665A 3.798<br>713A - 739A 3.370<br>717A - 566A 4.432<br>720A - 566A 4.461<br>720A - 568A 4.608<br>720A - 569A 3.700<br>720A - 570A 4.128<br>724A - 568A 4.297<br>724A - 570A 2.991<br>731A - 570A 4.684<br>731A - 626A 2.379<br>731A - 628A 4.708 | 115A - 523A 4.289<br>115A - 526A 2.777<br>116A - 501A 3.113<br>116A - 502A 4.539<br>116A - 503A 1.044<br>116A - 504A 4.592<br>116A - 520A 4.634<br>116A - 521A 3.327<br>117A - 501A 3.006<br>117A - 513A 2.789<br>117A - 518A 3.110<br>117A - 519A 3.943<br>117A - 520A 4.001<br>117A - 521A 4.623<br>118A - 501A 2.833<br>118A - 518A 3.987<br>118A - 519A 1.395<br>118A - 520A 3.141<br>118A - 521A 4.302<br>119A - 518A 4.399<br>119A - 519A 3.388<br>120A - 460A 2.542<br>120A - 517A 3.440<br>120A - 518A 4.820<br>120A - 519A 4.130<br>122A - 460A 3.962<br>383A - 504A 4.371<br>389A - 457A 4.136<br>389A - 460A 4.828<br>412A - 460A 2.752<br>414A - 456A 4.956<br>414A - 457A 4.839<br>414A - 460A 3.858<br>416A - 501A 2.611<br>416A - 503A 4.744<br>417A - 501A 4.708<br>418A - 503A 3.607<br>418A - 504A 1.782<br>419A - 504A 4.735<br>420A - 504A 4.718 |
| **Integrin β3** | 7A - 285A 3.835<br>10A - 285A 4.050<br>11A - 280A 4.653<br>11A - 282A 2.590<br>12A - 9A 4.110<br>12A - 12A 3.387<br>12A - 13A 2.649<br>12A - 16A 4.463<br>14A - 280A 3.354<br>14A - 287A 1.732<br>14A - 314A 4.993<br>15A - 12A 3.088<br>16A - 12A 4.517<br>18A - 259A 3.804<br>18A - 287A 3.088<br>19A - 8A 4.269<br>19A - 11A 3.664<br>19A - 12A 3.209<br>19A - 15A 4.106 | 52A - 33A 3.597<br>52A - 68A 2.934<br>53A - 68A 4.272<br>54A - 35A 2.680<br>54A - 67A 3.015<br>54A - 68A 2.416<br>55A - 67A 4.572<br>78A - 64A 4.325<br>78A - 65A 4.255<br>78A - 68A 2.868<br>78A - 69A 4.459<br>78A - 70A 4.654<br>79A - 70A 4.679<br>80A - 33A 3.532<br>80A - 70A 3.038<br>92A - 205A 4.528<br>117A - 204A 2.230<br>117A - 205A 3.859<br>189A - 238A 3.833 |

| | | |
|---|---|---|
| | 20A - 8A 4.953 | 189A - 239A 4.884 |
| | 22A - 259A 4.286 | 189A - 240A 4.754 |
| | 47A - 263A 4.681 | 190A - 238A 3.811 |
| | 47A - 265A 4.095 | 190A - 239A 4.228 |
| | 47A - 274A 2.187 | 193A - 238A 3.835 |
| | 59A - 880A 3.267 | 195A - 236A 3.396 |
| | 59A - 886A 4.413 | 195A - 238A 1.986 |
| | 60A - 886A 4.987 | 195A - 239A 4.801 |
| | 61A - 880A 4.522 | 196A - 320A 4.320 |
| | 61A - 886A 4.659 | 196A - 412A 3.497 |
| | 61A - 888A 4.425 | 196A - 413A 4.225 |
| | 68A - 263A 4.825 | 198A - 350A 3.753 |
| | 68A - 274A 4.767 | 201A - 238A 4.730 |
| | 69A - 236A 2.437 | 291A - 240A 4.843 |
| | 69A - 238A 4.642 | 292A - 230A 2.252 |
| | 69A - 239A 3.111 | 292A - 242A 4.486 |
| | 69A - 263A 4.593 | 293A - 230A 4.488 |
| | 70A - 233A 2.831 | 293A - 242A 4.590 |
| | 70A - 234A 3.835 | 294A - 229A 3.927 |
| | 70A - 241A 1.478 | 294A - 230A 3.997 |
| | 70A - 263A 4.776 | 294A - 242A 4.181 |
| | 73A - 236A 2.660 | 295A - 242A 4.353 |
| | 74A - 233A 4.596 | 295A - 244A 4.901 |
| | 74A - 234A 4.576 | 295A - 277A 4.644 |
| | 74A - 241A 4.927 | 295A - 279A 3.653 |
| | 79A - 236A 4.533 | 295A - 283A 3.103 |
| | 84A - 239A 4.375 | 295A - 284A 3.569 |
| | 84A - 263A 4.949 | 295A - 285A 2.529 |
| | 84A - 265A 2.414 | 298A - 279A 2.803 |
| | 84A - 274A 4.939 | 298A - 283A 4.927 |
| | 86A - 238A 4.445 | 300A - 256A 3.469 |
| | 86A - 267A 3.048 | 300A - 278A 3.380 |
| | 87A - 267A 1.885 | 300A - 279A 2.986 |
| | 87A - 272A 4.895 | 300A - 280A 2.507 |
| | 88A - 238A 4.182 | 302A - 277A 3.634 |
| | 88A - 267A 3.643 | 302A - 278A 4.350 |
| | 88A - 268A 3.091 | 302A - 279A 3.749 |
| | 89A - 268A 4.196 | 302A - 283A 4.804 |
| | 89A - 269A 3.950 | 311A - 276A 3.626 |
| | 89A - 270A 4.033 | 311A - 277A 4.016 |
| | 90A - 268A 1.988 | 311A - 285A 3.159 |
| | 90A - 269A 4.038 | 313A - 242A 4.754 |
| | 91A - 269A 2.346 | 313A - 287A 3.147 |
| | 124A - 272A 3.976 | 314A - 242A 3.283 |
| | 125A - 272A 2.853 | 314A - 285A 3.391 |
| | 126A - 267A 2.792 | 314A - 286A 3.672 |
| | 126A - 270A 3.690 | 314A - 287A 2.270 |
| | 126A - 272A 2.762 | 437A - 160A 3.453 |
| | 127A - 270A 2.114 | 437A - 161A 2.002 |
| | 127A - 295A 3.944 | 437A - 162A 4.497 |
| | 127A - 302A 4.133 | 438A - 197A 4.435 |
| | 128A - 270A 3.767 | 438A - 199A 4.957 |
| | 129A - 270A 4.693 | 438A - 200A 2.475 |
| | 129A - 297A 3.501 | 438A - 201A 4.656 |
| | 129A - 300A 4.735 | 438A - 202A 3.408 |
| | 405A - 334A 4.498 | 439A - 195A 2.591 |
| | 405A - 348A 4.379 | 439A - 196A 4.412 |
| | 406A - 348A 3.439 | 439A - 202A 3.100 |
| | 406A - 373A 4.244 | 439A - 203A 3.140 |
| | 407A - 334A 2.537 | 439A - 204A 3.490 |

| | | |
|---|---|---|
| | 407A - 348A 2.770 | 439A - 206A 3.428 |
| | 426A - 302A 3.972 | 439A - 208A 4.362 |
| | 509A - 873A 4.315 | 439A - 209A 3.723 |
| | 512A - 914A 4.460 | 441A - 209A 4.384 |
| | 607A - 352A 3.212 | 454A - 208A 3.723 |
| | 607A - 354A 4.576 | 456A - 204A 4.030 |
| | 607A - 369A 2.767 | 457A - 204A 4.915 |
| | 607A - 370A 4.960 | 458A - 202A 3.004 |
| | 607A - 371A 4.561 | 458A - 203A 4.839 |
| | 651A - 397A 3.312 | 458A - 204A 1.849 |
| | 654A - 391A 3.416 | 462A - 199A 3.867 |
| | 654A - 399A 4.312 | 462A - 200A 4.020 |
| | 654A - 401A 4.743 | 467A - 199A 4.023 |
| | 655A - 391A 3.206 | 468A - 25A 3.840 |
| | 655A - 392A 4.740 | 468A - 31A 4.951 |
| | 655A - 393A 2.217 | 468A - 32A 2.907 |
| | 655A - 398A 4.963 | 468A - 33A 1.793 |
| | 655A - 399A 2.059 | 468A - 34A 3.953 |
| | 656A - 391A 2.373 | 470A - 23A 4.770 |
| | 657A - 393A 3.790 | 483A - 68A 3.127 |
| | | 486A - 68A 4.371 |
| **Integrin β4** | 6A - 287A 3.163 | 172A - 301A 4.053 |
| | 7A - 287A 3.604 | 172A - 306A 4.342 |
| | 7A - 310A 2.856 | 177A - 264A 3.873 |
| | 7A - 311A 3.407 | 177A - 265A 2.109 |
| | 7A - 312A 4.509 | 177A - 266A 4.303 |
| | 7A - 314A 4.144 | 178A - 265A 3.379 |
| | 10A - 280A 1.833 | 178A - 267A 4.892 |
| | 10A - 282A 4.195 | 184A - 265A 3.597 |
| | 10A - 287A 1.830 | 184A - 266A 4.735 |
| | 11A - 280A 4.365 | 184A - 297A 4.419 |
| | 12A - 12A 4.813 | 186A - 297A 4.978 |
| | 13A - 12A 4.388 | 186A - 312A 3.688 |
| | 13A - 15A 3.950 | 186A - 314A 3.443 |
| | 13A - 16A 4.633 | 189A - 297A 2.987 |
| | 13A - 245A 3.415 | 189A - 299A 4.255 |
| | 13A - 247A 3.623 | 189A - 312A 3.560 |
| | 13A - 257A 3.400 | 190A - 309A 3.111 |
| | 13A - 282A 3.901 | 190A - 310A 4.318 |
| | 14A - 280A 2.747 | 190A - 311A 4.751 |
| | 14A - 281A 4.679 | 190A - 312A 3.145 |
| | 14A - 282A 2.157 | 191A - 309A 4.795 |
| | 14A - 285A 4.544 | 196A - 309A 3.258 |
| | 16A - 12A 2.895 | 197A - 301A 2.469 |
| | 16A - 13A 4.274 | 197A - 306A 3.456 |
| | 16A - 16A 3.087 | 197A - 307A 4.017 |
| | 17A - 247A 4.336 | 197A - 308A 3.212 |
| | 17A - 255A 4.767 | 197A - 309A 4.190 |
| | 17A - 282A 2.653 | 198A - 299A 3.681 |
| | 20A - 20A 4.394 | 198A - 301A 4.610 |
| | 23A - 100A 2.709 | 198A - 308A 4.051 |
| | 23A - 101A 3.202 | 198A - 309A 2.007 |
| | 23A - 102A 2.512 | 198A - 310A 3.905 |
| | 23A - 103A 4.910 | 199A - 263A 4.838 |
| | 25A - 98A 4.499 | 203A - 263A 4.908 |
| | 25A - 100A 4.202 | 203A - 301A 2.409 |
| | 57A - 93A 3.392 | 218A - 306A 3.700 |
| | 57A - 95A 2.627 | 221A - 218A 4.833 |
| | 58A - 95A 4.758 | 221A - 219A 3.879 |

| | | |
|---|---|---|
| | 58A - 96A 4.145<br>58A - 98A 4.020<br>58A - 119A 2.823<br>68A - 106A 4.723<br>69A - 106A 2.836<br>70A - 98A 2.337<br>70A - 106A 4.939<br>71A - 119A 3.767<br>71A - 123A 4.955<br>467A - 47A 2.657<br>468A - 45A 4.971<br>468A - 47A 4.616<br>468A - 49A 4.773<br>470A - 45A 2.907<br>470A - 87A 3.204<br>472A - 45A 4.606<br>486A - 45A 3.447<br>488A - 92A 4.650<br>489A - 44A 4.451<br>489A - 92A 3.251<br>490A - 44A 2.712<br>490A - 45A 4.219<br>491A - 45A 4.059<br>492A - 43A 3.683<br>492A - 44A 3.207<br>492A - 45A 2.370<br>493A - 45A 3.936<br>500A - 40A 3.987<br>502A - 38A 4.869<br>503A - 38A 4.304<br>503A - 39A 2.939<br>503A - 40A 2.602<br>504A - 39A 3.935<br>504A - 40A 2.827<br>504A - 41A 4.372<br>504A - 42A 3.829<br>506A - 39A 2.627<br>507A - 39A 4.686<br>508A - 39A 4.413<br>522A - 42A 2.102<br>522A - 92A 4.237<br>523A - 42A 3.606<br>524A - 42A 3.973<br>524A - 92A 3.897<br>537A - 39A 4.248<br>999A - 497A 3.425<br>999A - 498A 4.442<br>1013A - 498A 4.210<br>1013A - 504A 4.993<br>1015A - 497A 4.571<br>1015A - 498A 3.607<br>1053A - 537A 2.284<br>1054A - 537A 3.918<br>1055A - 484A 4.724<br>1055A - 529A 4.819<br>1056A - 482A 3.463<br>1056A - 484A 3.626<br>1056A - 529A 4.801<br>1056A - 531A 2.432<br>1056A - 537A 3.025 | 221A - 304A 2.944<br>221A - 305A 4.864<br>221A - 306A 2.167<br>221A - 307A 3.266<br>222A - 301A 4.912<br>222A - 306A 2.567<br>223A - 306A 3.743<br>223A - 307A 2.696<br>223A - 308A 3.519<br>223A - 309A 3.295<br>277A - 267A 4.035<br>278A - 267A 3.662<br>279A - 267A 3.872<br>290A - 274A 2.677<br>290A - 275A 4.456<br>291A - 275A 4.678<br>291A - 303A 4.587<br>292A - 259A 3.230<br>292A - 261A 3.713<br>292A - 274A 4.205<br>292A - 275A 1.860<br>292A - 303A 2.429<br>294A - 261A 2.259<br>296A - 274A 4.304<br>298A - 264A 4.975<br>299A - 264A 3.050<br>299A - 265A 3.620<br>299A - 266A 2.173<br>299A - 267A 3.030<br>299A - 269A 4.876<br>300A - 269A 4.717<br>300A - 270A 3.284<br>1643A - 357A 3.210<br>1643A - 383A 4.879<br>1643A - 384A 4.650<br>1643A - 385A 3.102<br>1645A - 357A 4.503<br>1645A - 385A 3.334<br>1647A - 354A 4.889<br>1655A - 349A 4.786<br>1655A - 353A 2.318<br>1655A - 354A 2.959<br>1655A - 355A 4.918<br>1657A - 354A 4.736<br>1657A - 356A 4.078<br>1658A - 356A 3.323<br>1658A - 357A 3.127<br>1659A - 357A 2.380<br>1659A - 382A 4.357<br>1659A - 383A 4.830<br>1660A - 357A 4.298<br>1660A - 359A 4.138<br>1688A - 295A 3.238<br>1688A - 316A 4.437<br>1689A - 293A 3.367<br>1689A - 294A 3.474<br>1689A - 295A 3.658<br>1689A - 317A 3.264<br>1691A - 358A 4.651<br>1692A - 292A 3.259 |

| | | |
|---|---|---|
| | 1057A - 484A 2.667<br>1057A - 486A 4.557<br>1058A - 484A 3.321<br>1058A - 497A 2.899<br>1058A - 498A 4.461<br>1058A - 500A 4.091<br>1058A - 501A 1.394<br>1058A - 502A 4.404<br>1059A - 482A 2.897<br>1059A - 501A 4.438<br>1059A - 531A 4.110<br>1060A - 497A 4.624<br>1060A - 498A 4.305<br>1060A - 501A 2.199<br>1060A - 502A 3.336<br>1060A - 503A 4.203<br>1062A - 503A 3.485<br>1062A - 504A 4.849 | 1692A - 293A 2.781<br>1692A - 294A 2.924<br>1692A - 295A 4.281<br>1693A - 356A 4.293<br>1693A - 357A 3.027<br>1693A - 358A 4.104<br>1693A - 359A 3.910<br>1694A - 292A 2.733<br>1694A - 354A 4.733<br>1695A - 292A 2.957<br>1696A - 292A 3.040<br>1696A - 351A 3.165<br>1696A - 353A 4.703<br>1760A - 354A 3.705<br>1760A - 385A 3.503<br>1760A - 386A 2.866<br>1760A - 387A 4.587<br>1761A - 385A 2.644<br>1761A - 386A 2.738<br>1762A - 384A 3.182<br>1762A - 385A 4.020<br>1762A - 386A 1.251<br>1762A - 387A 4.565<br>1762A - 388A 4.764<br>1763A - 385A 4.939<br>1768A - 346A 4.095<br>1768A - 354A 4.733<br>1768A - 386A 4.184<br>1846A - 385A 2.535 |
| **Integrin β5** | 24A - 238A 3.126<br>28A - 268A 4.102<br>29A - 269A 4.374<br>30A - 269A 4.696<br>31A - 269A 3.034<br>32A - 269A 4.467<br>47A - 269A 2.632<br>48A - 297A 3.986<br>49A - 269A 3.116<br>70A - 269A 2.610<br>71A - 299A 3.201<br>72A - 298A 2.020<br>72A - 427A 2.957<br>72A - 428A 3.637<br>72A - 429A 4.658<br>81A - 339A 4.258<br>81A - 340A 3.642<br>81A - 342A 4.915<br>82A - 340A 3.867<br>82A - 341A 4.990<br>84A - 339A 4.547<br>84A - 340A 3.675<br>86A - 299A 3.686<br>86A - 339A 2.507<br>87A - 339A 4.715<br>88A - 297A 2.469<br>88A - 298A 3.167<br>88A - 299A 3.986<br>88A - 337A 4.180<br>88A - 339A 3.694 | 193A - 295A 1.789<br>193A - 296A 4.954<br>193A - 314A 3.277<br>193A - 316A 4.417<br>194A - 295A 3.106<br>194A - 316A 3.627<br>196A - 314A 3.756<br>197A - 314A 4.536<br>198A - 297A 4.372<br>198A - 312A 4.980<br>198A - 314A 3.349<br>199A - 312A 2.606<br>199A - 313A 4.036<br>199A - 314A 4.856<br>200A - 312A 2.880<br>201A - 226A 3.932<br>201A - 311A 4.065<br>201A - 312A 3.663<br>202A - 312A 4.970<br>248A - 231A 3.959<br>248A - 295A 4.282<br>248A - 314A 2.407<br>248A - 316A 2.755<br>290A - 357A 3.727<br>293A - 355A 4.975<br>293A - 356A 3.609<br>293A - 357A 4.250<br>293A - 358A 1.869<br>294A - 294A 4.420<br>294A - 295A 4.224 |

| | | |
|---|---|---|
| | 89A - 336A 4.747 | 529A - 504A 4.555 |
| | 91A - 334A 4.247 | 530A - 504A 2.933 |
| | 91A - 348A 3.219 | 541A - 504A 4.026 |
| | 93A - 348A 3.523 | 543A - 503A 4.826 |
| | 93A - 373A 3.112 | 545A - 500A 4.808 |
| | 93A - 375A 4.226 | 545A - 501A 2.974 |
| | 96A - 393A 2.981 | 545A - 502A 2.934 |
| | 97A - 393A 3.969 | 545A - 503A 3.893 |
| | 98A - 393A 2.009 | 548A - 504A 1.804 |
| | 98A - 394A 3.612 | 558A - 501A 3.042 |
| | 98A - 396A 4.704 | 558A - 503A 4.009 |
| | 98A - 397A 2.731 | 558A - 504A 4.095 |
| | 98A - 398A 4.013 | 558A - 519A 3.628 |
| | 99A - 371A 1.567 | 559A - 513A 4.866 |
| | 99A - 393A 4.606 | 560A - 518A 4.281 |
| | 99A - 394A 4.762 | 561A - 516A 4.152 |
| | 99A - 395A 4.233 | 561A - 518A 3.501 |
| | 99A - 396A 3.236 | 562A - 516A 2.200 |
| | 99A - 397A 3.523 | 562A - 517A 3.494 |
| | 100A - 397A 2.803 | 562A - 518A 2.842 |
| | 101A - 367A 2.692 | 562A - 519A 4.326 |
| | 101A - 396A 3.702 | 563A - 463A 3.762 |
| | 106A - 397A 4.212 | 563A - 467A 4.760 |
| | 110A - 397A 2.598 | 563A - 470A 4.998 |
| | 111A - 397A 4.270 | 563A - 471A 3.945 |
| | 112A - 397A 4.804 | 563A - 516A 3.184 |
| | 115A - 371A 3.980 | 564A - 463A 4.521 |
| | 115A - 373A 3.637 | 564A - 469A 4.235 |
| | 115A - 393A 4.673 | 564A - 516A 4.276 |
| | 116A - 373A 3.511 | 565A - 460A 4.205 |
| | 127A - 297A 2.494 | 587A - 472A 4.755 |
| | 127A - 300A 4.755 | 588A - 472A 2.698 |
| | 130A - 334A 3.960 | 588A - 515A 3.940 |
| | 130A - 348A 3.838 | 588A - 516A 4.972 |
| | 131A - 348A 3.776 | 589A - 472A 4.744 |
| | 132A - 348A 4.347 | 589A - 515A 4.561 |
| | 132A - 350A 2.327 | 589A - 516A 2.719 |
| | 132A - 351A 3.669 | 591A - 470A 4.040 |
| | 132A - 372A 3.741 | 591A - 471A 1.895 |
| | 132A - 373A 3.792 | 591A - 472A 2.592 |
| | 134A - 350A 4.700 | 592A - 471A 4.719 |
| | 134A - 351A 3.463 | 614A - 470A 3.056 |
| | 134A - 352A 4.906 | 615A - 470A 2.613 |
| | 134A - 371A 0.887 | 615A - 471A 4.161 |
| | 134A - 372A 3.000 | 616A - 470A 4.661 |
| | 134A - 373A 3.571 | 617A - 469A 2.790 |
| | 134A - 395A 4.978 | 617A - 470A 2.531 |
| | 136A - 367A 2.457 | |
| | 136A - 368A 3.304 | |
| | 136A - 371A 4.831 | |
| | 136A - 395A 3.889 | |
| | 136A - 396A 4.338 | |
| | 405A - 330A 3.005 | |
| | 405A - 352A 4.842 | |
| | 406A - 354A 4.306 | |
| | 407A - 354A 2.669 | |
| | 408A - 354A 4.276 | |
| | 409A - 306A 3.032 | |
| | 409A - 330A 2.798 | |
| | 409A - 354A 4.030 | |

| | | |
|---|---|---|
| | 410A - 306A 3.851<br>410A - 330A 4.771<br>411A - 306A 2.883<br>411A - 330A 4.167<br>422A - 356A 4.892<br>422A - 368A 3.313<br>423A - 368A 2.315<br>424A - 368A 3.530<br>424A - 369A 4.674<br>425A - 368A 3.406<br>425A - 369A 2.585<br>426A - 352A 3.185<br>426A - 368A 4.194<br>426A - 369A 3.033<br>426A - 370A 4.586<br>426A - 371A 2.282<br>426A - 395A 4.076<br>427A - 352A 3.982<br>428A - 352A 3.913<br>430A - 332A 4.465 | |
| **Integrin β6** | 10A - 501A 4.742<br>10A - 502A 3.469<br>10A - 503A 3.996<br>13A - 480A 4.746<br>13A - 481A 2.417<br>13A - 482A 3.485<br>13A - 503A 4.263<br>13A - 532A 4.708<br>13A - 533A 3.087<br>14A - 482A 3.672<br>14A - 483A 4.858<br>14A - 484A 3.891<br>14A - 501A 3.814<br>14A - 502A 4.802<br>17A - 531A 3.739<br>17A - 535A 4.898<br>17A - 537A 2.361<br>37A - 539A 4.837<br>38A - 529A 4.068<br>38A - 537A 2.324<br>38A - 539A 4.425<br>77A - 541A 4.389<br>78A - 541A 3.421<br>113A - 539A 4.859<br>114A - 452A 3.663<br>114A - 453A 3.800<br>114A - 454A 4.387<br>114A - 537A 4.079<br>114A - 538A 3.076<br>114A - 539A 3.243<br>115A - 539A 4.593<br>115A - 541A 4.188<br>116A - 453A 4.616<br>116A - 454A 2.170<br>116A - 455A 2.155<br>116A - 456A 4.474<br>116A - 458A 4.651<br>116A - 539A 4.209<br>116A - 540A 4.769 | 1A - 126A 4.629<br>3A - 126A 4.545<br>3A - 191A 4.883<br>3A - 213A 2.304<br>6A - 256A 2.242<br>6A - 257A 2.930<br>6A - 258A 4.002<br>7A - 256A 4.172<br>9A - 277A 2.837<br>10A - 256A 3.072<br>10A - 278A 4.621<br>10A - 279A 4.071<br>10A - 280A 4.216<br>13A - 273A 4.731<br>13A - 275A 3.610<br>13A - 276A 3.187<br>13A - 277A 3.322<br>13A - 285A 4.763<br>178A - 470A 4.315<br>179A - 470A 3.045<br>180A - 470A 4.917<br>274A - 340A 2.115<br>274A - 374A 3.518<br>274A - 392A 4.140<br>274A - 394A 4.073<br>277A - 340A 4.576<br>277A - 469A 3.846<br>278A - 469A 4.373<br>278A - 470A 4.264<br>279A - 469A 2.415<br>279A - 470A 2.868<br>280A - 468A 4.919<br>280A - 469A 1.468<br>280A - 470A 4.432<br>281A - 469A 3.748<br>282A - 337A 4.165<br>282A - 338A 3.355<br>282A - 394A 2.612<br>283A - 337A 4.542 |

| | | |
|---|---|---|
| | 116A - 541A 4.102 | 283A - 394A 4.663 |
| | 117A - 458A 3.131 | 284A - 372A 4.854 |
| | 118A - 456A 4.821 | 284A - 373A 3.966 |
| | 118A - 457A 2.839 | 284A - 374A 2.704 |
| | 118A - 458A 2.691 | 284A - 394A 2.504 |
| | 180A - 267A 4.492 | 286A - 376A 3.659 |
| | 180A - 270A 3.346 | 287A - 376A 4.029 |
| | 180A - 272A 4.916 | 304A - 374A 3.044 |
| | 181A - 272A 4.848 | 304A - 394A 2.902 |
| | 232A - 267A 2.964 | 307A - 374A 3.741 |
| | 233A - 267A 4.630 | 307A - 375A 2.191 |
| | 233A - 272A 2.335 | 307A - 376A 4.617 |
| | 234A - 265A 2.991 | 307A - 379A 4.153 |
| | 234A - 274A 3.307 | 308A - 379A 4.972 |
| | 235A - 272A 4.937 | 308A - 380A 4.959 |
| | 235A - 274A 4.344 | 309A - 376A 4.867 |
| | 269A - 274A 4.109 | 309A - 378A 4.621 |
| | 269A - 291A 4.484 | 309A - 379A 3.424 |
| | 272A - 291A 4.123 | 309A - 380A 4.381 |
| | 272A - 292A 4.743 | 310A - 374A 3.701 |
| | 272A - 293A 4.137 | 310A - 375A 4.460 |
| | 272A - 304A 3.596 | 310A - 376A 3.215 |
| | 272A - 305A 4.562 | 313A - 376A 2.389 |
| | 272A - 306A 1.346 | 314A - 376A 3.721 |
| | 273A - 306A 4.751 | 340A - 359A 3.910 |
| | 274A - 332A 3.182 | 340A - 380A 3.986 |
| | 275A - 293A 3.598 | 340A - 382A 4.482 |
| | 275A - 295A 4.889 | 380A - 406A 3.952 |
| | 275A - 304A 3.981 | 380A - 411A 4.302 |
| | 277A - 302A 3.787 | 381A - 411A 4.095 |
| | 277A - 334A 3.764 | 383A - 236A 4.181 |
| | 278A - 270A 3.416 | 383A - 239A 4.177 |
| | 278A - 271A 4.763 | 385A - 233A 3.944 |
| | 278A - 272A 3.391 | 397A - 228A 3.549 |
| | 278A - 294A 4.535 | 397A - 229A 3.904 |
| | 278A - 295A 1.375 | 397A - 230A 4.200 |
| | 278A - 296A 3.952 | 397A - 231A 3.421 |
| | 278A - 302A 3.160 | 397A - 232A 3.960 |
| | 307A - 306A 4.803 | 398A - 231A 3.681 |
| | 307A - 330A 4.609 | 398A - 232A 3.306 |
| | 308A - 306A 3.757 | 399A - 232A 4.650 |
| | 332A - 276A 2.760 | 420A - 236A 4.805 |
| | 333A - 275A 4.940 | 544A - 291A 4.949 |
| | 333A - 276A 3.469 | 544A - 350A 3.735 |
| | 333A - 291A 4.387 | 544A - 351A 2.792 |
| | 336A - 276A 2.814 | 544A - 352A 4.877 |
| | 336A - 289A 3.148 | 546A - 236A 4.814 |
| | 336A - 290A 4.936 | 546A - 237A 3.212 |
| | 336A - 291A 3.846 | 546A - 238A 4.904 |
| | 336A - 308A 3.683 | 551A - 270A 4.343 |
| | 336A - 310A 3.752 | 565A - 270A 3.772 |
| | 337A - 291A 2.846 | 577A - 261A 2.773 |
| | 339A - 308A 4.406 | 577A - 262A 1.458 |
| | 339A - 309A 4.653 | 577A - 263A 3.496 |
| | 339A - 310A 1.612 | 577A - 264A 2.115 |
| | 339A - 328A 3.516 | 577A - 265A 3.408 |
| | 340A - 307A 4.017 | 577A - 266A 4.926 |
| | 340A - 308A 2.238 | 577A - 271A 3.587 |
| | 340A - 309A 4.809 | 577A - 273A 4.669 |
| | 340A - 328A 3.103 | 577A - 274A 3.110 |

| | | |
|---|---|---|
| | 380A - 368A 3.571 | 577A - 275A 4.994 |
| | 383A - 357A 4.788 | 578A - 264A 3.342 |
| | 383A - 448A 2.752 | 578A - 265A 2.209 |
| | 384A - 453A 4.967 | 578A - 266A 1.960 |
| | 385A - 450A 4.497 | 578A - 267A 3.460 |
| | 385A - 453A 1.732 | 578A - 269A 4.919 |
| | 385A - 475A 4.093 | 579A - 265A 4.182 |
| | 386A - 453A 3.053 | 579A - 266A 4.709 |
| | 386A - 475A 3.198 | 579A - 267A 4.892 |
| | 387A - 453A 4.377 | 580A - 264A 4.962 |
| | 387A - 473A 2.690 | 580A - 265A 3.367 |
| | 387A - 475A 2.001 | 596A - 266A 4.495 |
| | 389A - 473A 4.299 | 596A - 267A 3.145 |
| | 395A - 473A 3.389 | 597A - 267A 3.547 |
| | 396A - 475A 3.937 | 597A - 268A 4.768 |
| | 397A - 475A 2.143 | 597A - 351A 3.719 |
| | 398A - 475A 2.758 | 598A - 267A 4.376 |
| | 399A - 475A 4.925 | 608A - 267A 3.950 |
| | 414A - 453A 3.963 | 608A - 292A 2.576 |
| | 414A - 455A 2.569 | 608A - 294A 3.873 |
| | 414A - 456A 4.655 | 609A - 264A 4.654 |
| | 414A - 473A 3.420 | 609A - 265A 1.672 |
| | 415A - 453A 3.971 | 609A - 266A 2.440 |
| | 416A - 453A 1.599 | 609A - 267A 4.851 |
| | 416A - 454A 3.687 | |
| | 416A - 455A 4.733 | |
| | 420A - 448A 3.971 | |
| | 544A - 317A 3.075 | |
| | 544A - 318A 1.585 | |
| | 544A - 319A 3.452 | |
| | 544A - 364A 4.612 | |
| | 546A - 320A 4.635 | |
| | 551A - 320A 2.351 | |
| | 565A - 320A 4.182 | |
| | 597A - 317A 3.396 | |
| | 607A - 285A 4.162 | |
| | 608A - 283A 4.325 | |
| | 608A - 284A 4.802 | |
| | 608A - 285A 4.100 | |
| | 612A - 280A 3.540 | |
| | 612A - 282A 2.759 | |
| | 612A - 283A 4.915 | |
| | 612A - 285A 2.365 | |
| | 615A - 102A 4.457 | |
| | 617A - 102A 4.569 | |
| | 618A - 13A 3.224 | |
| | 655A - 1A 4.328 | |
| | 657A - 3A 3.800 | |
| | 659A - 3A 2.657 | |
| | 664A - 2A 4.555 | |
| | 664A - 3A 1.540 | |
| | 664A - 4A 3.289 | |
| | 664A - 5A 3.987 | |
| | 666A - 3A 4.933 | |
| | 666A - 5A 4.556 | |
| | 676A - 2A 3.366 | |
| | 678A - 3A 3.954 | |
| | 690A - 1A 2.418 | |
| | 690A - 2A 4.932 | |
| | 692A - 1A 3.559 | |

| | | |
|---|---|---|
| | 692A - 2A 3.457<br>694A - 2A 3.323 | |
| **Integrin β7** | 6A - 299A 4.968<br>6A - 336A 3.443<br>6A - 337A 3.149<br>6A - 338A 2.605<br>6A - 339A 4.484<br>6A - 346A 4.648<br>7A - 297A 3.358<br>9A - 338A 4.784<br>9A - 339A 2.048<br>9A - 340A 4.470<br>10A - 297A 2.226<br>10A - 298A 3.001<br>10A - 299A 3.375<br>10A - 300A 3.805<br>10A - 337A 4.097<br>10A - 339A 3.700<br>11A - 297A 3.037<br>13A - 299A 3.020<br>13A - 339A 3.777<br>14A - 269A 2.953<br>14A - 297A 3.094<br>14A - 298A 4.177<br>15A - 269A 3.507<br>17A - 268A 4.044<br>17A - 269A 3.161<br>17A - 298A 3.948<br>18A - 269A 3.251<br>19A - 269A 4.890<br>183A - 314A 4.784<br>197A - 475A 4.852<br>198A - 473A 2.644<br>200A - 457A 4.689<br>200A - 473A 4.676<br>235A - 312A 4.077<br>236A - 310A 4.607<br>236A - 312A 1.616<br>237A - 312A 2.811<br>238A - 309A 4.367<br>238A - 310A 2.444<br>238A - 311A 4.012<br>238A - 312A 4.178<br>271A - 308A 3.747<br>271A - 309A 3.824<br>271A - 328A 3.534<br>272A - 310A 2.918<br>273A - 310A 3.211<br>274A - 276A 4.635<br>274A - 277A 4.396<br>274A - 278A 4.267<br>274A - 289A 3.077<br>274A - 310A 3.349<br>275A - 278A 4.676<br>276A - 287A 2.963<br>277A - 287A 3.102<br>277A - 310A 3.049<br>279A - 287A 4.781 | 1A - 535A 3.822<br>4A - 451A 4.064<br>5A - 450A 4.427<br>5A - 451A 3.541<br>5A - 534A 4.693<br>5A - 535A 4.348<br>8A - 451A 2.017<br>8A - 534A 3.311<br>9A - 451A 4.511<br>9A - 534A 2.352<br>10A - 534A 4.673<br>12A - 427A 3.852<br>12A - 428A 4.886<br>12A - 429A 4.125<br>12A - 430A 0.794<br>12A - 431A 4.133<br>12A - 451A 4.258<br>12A - 534A 2.558<br>13A - 423A 3.959<br>13A - 427A 4.501<br>13A - 534A 4.364<br>509A - 232A 3.990<br>509A - 318A 4.981<br>510A - 318A 3.841<br>510A - 319A 3.867<br>512A - 361A 2.109<br>512A - 363A 4.531<br>512A - 378A 3.990<br>512A - 380A 3.780<br>512A - 408A 3.620<br>515A - 361A 4.947<br>515A - 380A 4.152<br>517A - 378A 2.963<br>517A - 379A 4.802<br>517A - 380A 4.474<br>537A - 380A 4.514<br>539A - 375A 4.542<br>539A - 376A 3.118<br>539A - 378A 4.643<br>539A - 379A 4.569<br>541A - 374A 2.426<br>541A - 375A 2.943<br>541A - 376A 2.773<br>542A - 374A 4.630<br>542A - 375A 2.541<br>542A - 379A 2.438<br>542A - 380A 4.624<br>542A - 381A 4.918<br>543A - 374A 4.476<br>543A - 375A 4.270<br>544A - 374A 4.785<br>545A - 338A 4.976<br>545A - 340A 4.054<br>545A - 372A 3.768<br>545A - 373A 2.399<br>545A - 374A 2.413 |

| | | |
|---|---|---|
| | 280A - 280A 3.226 | 545A - 375A 4.869 |
| | 280A - 285A 4.702 | 545A - 394A 2.326 |
| | 280A - 287A 2.840 | 546A - 394A 4.130 |
| | 280A - 314A 3.783 | 546A - 468A 4.885 |
| | 295A - 766A 4.631 | 546A - 469A 2.928 |
| | 295A - 837A 3.963 | 547A - 337A 4.196 |
| | 295A - 838A 4.522 | 547A - 338A 3.216 |
| | 296A - 766A 3.457 | 547A - 394A 4.868 |
| | 309A - 276A 2.943 | 547A - 467A 4.337 |
| | 334A - 308A 4.314 | 547A - 468A 4.312 |
| | 337A - 291A 3.222 | 547A - 469A 4.403 |
| | 337A - 306A 4.737 | 548A - 337A 2.526 |
| | 338A - 291A 4.567 | 548A - 338A 4.428 |
| | 340A - 274A 4.909 | 548A - 394A 4.120 |
| | 341A - 274A 3.378 | 552A - 469A 3.227 |
| | 341A - 275A 4.371 | 552A - 470A 4.011 |
| | 341A - 276A 3.620 | 554A - 470A 3.536 |
| | 341A - 291A 4.218 | 568A - 470A 2.478 |
| | 384A - 238A 4.852 | 574A - 469A 3.809 |
| | 401A - 267A 4.556 | 574A - 470A 4.767 |
| | 402A - 236A 4.867 | 576A - 470A 3.512 |
| | 402A - 238A 2.350 | 581A - 470A 4.247 |
| | 402A - 239A 3.374 | 581A - 471A 3.219 |
| | 402A - 265A 4.027 | 581A - 472A 4.446 |
| | 402A - 266A 4.703 | 581A - 516A 4.849 |
| | | 597A - 469A 4.565 |
| | | 598A - 470A 4.261 |
| | | 598A - 471A 4.251 |
| | | 599A - 469A 4.894 |
| | | 607A - 460A 3.350 |
| | | 608A - 460A 3.926 |
| | | 609A - 460A 3.981 |
| | | 609A - 463A 4.411 |
| | | 609A - 516A 3.172 |
| | | 609A - 517A 1.955 |
| | | 609A - 518A 2.911 |
| | | 610A - 518A 4.062 |
| | | 613A - 501A 3.096 |
| | | 613A - 519A 3.410 |
| | | 656A - 502A 3.339 |
| | | 668A - 504A 1.946 |
| **Integrin β8** | 225A - 312A 3.778 | 630A - 285A 2.874 |
| | 261A - 839A 3.020 | 640A - 240A 4.155 |
| | 261A - 862A 2.722 | 640A - 287A 4.602 |
| | 262A - 765A 4.946 | 641A - 230A 4.415 |
| | 262A - 766A 4.018 | 641A - 240A 4.824 |
| | 262A - 767A 4.742 | 641A - 241A 4.687 |
| | 262A - 839A 4.618 | 641A - 242A 3.017 |
| | 264A - 839A 3.832 | 641A - 286A 4.869 |
| | 264A - 840A 4.484 | 641A - 287A 1.923 |
| | 265A - 765A 3.283 | 643A - 230A 4.970 |
| | 265A - 766A 1.817 | 643A - 233A 4.442 |
| | 265A - 767A 3.910 | 643A - 240A 4.894 |
| | 265A - 768A 4.912 | 644A - 233A 3.643 |
| | 265A - 837A 4.771 | 645A - 231A 4.957 |
| | 265A - 838A 4.499 | 645A - 232A 3.490 |
| | 265A - 839A 2.810 | 645A - 233A 2.775 |
| | 268A - 838A 4.185 | 645A - 234A 4.075 |
| | 268A - 839A 2.731 | 647A - 231A 3.512 |

| | | |
|---|---|---|
| | 268A - 840A 3.254<br>268A - 841A 4.375<br>269A - 766A 3.040<br>269A - 837A 2.889<br>269A - 838A 4.883<br>277A - 310A 3.940<br>278A - 310A 3.394<br>279A - 287A 4.956<br>279A - 310A 3.191<br>279A - 311A 4.634<br>313A - 287A 3.686<br>314A - 314A 4.629<br>315A - 310A 4.526<br>315A - 312A 2.857<br>316A - 314A 1.949<br>317A - 313A 3.679<br>317A - 314A 4.488<br>318A - 312A 4.901<br>318A - 313A 2.867<br>318A - 314A 4.175<br>318A - 315A 2.429<br>318A - 323A 2.772<br>318A - 447A 3.392<br>318A - 449A 2.787<br>319A - 311A 4.321<br>319A - 312A 2.024<br>319A - 313A 2.431<br>319A - 314A 3.116<br>319A - 323A 4.855<br>320A - 312A 4.181<br>321A - 313A 4.209<br>321A - 323A 2.073<br>321A - 447A 1.396<br>321A - 448A 2.396<br>321A - 449A 2.962<br>321A - 450A 2.946<br>322A - 311A 4.139<br>322A - 312A 2.907<br>322A - 313A 2.060<br>322A - 325A 2.208<br>322A - 447A 3.901<br>324A - 475A 4.126<br>325A - 475A 4.474<br>326A - 450A 3.585<br>326A - 453A 4.806<br>326A - 475A 2.495<br>326A - 476A 3.893<br>327A - 473A 4.423<br>327A - 474A 4.026<br>327A - 475A 3.088<br>327A - 476A 4.651<br>328A - 475A 4.458<br>328A - 476A 1.954<br>328A - 477A 3.811<br>371A - 6A 3.095<br>371A - 8A 3.537<br>371A - 9A 2.823<br>371A - 12A 4.100<br>372A - 12A 4.901<br>373A - 287A 3.621 | 647A - 232A 1.883<br>647A - 233A 3.682<br>682A - 377A 4.132<br>682A - 378A 4.428<br>684A - 377A 2.425<br>684A - 378A 2.686<br>689A - 411A 4.684<br>691A - 363A 3.248<br>691A - 406A 4.132<br>691A - 408A 3.185<br>692A - 408A 4.710<br>693A - 361A 3.806<br>703A - 377A 4.045<br>703A - 378A 2.391<br>703A - 379A 4.498<br>737A - 469A 4.399<br>740A - 470A 3.100<br>741A - 469A 3.764<br>741A - 470A 3.706<br>744A - 470A 3.257<br>744A - 471A 3.493<br>745A - 469A 4.880<br>745A - 470A 4.376<br>745A - 471A 4.800<br>748A - 470A 4.802<br>748A - 471A 2.981<br>748A - 472A 3.942<br>748A - 473A 4.894<br>748A - 515A 3.814<br>748A - 516A 3.866<br>751A - 472A 4.065<br>751A - 515A 3.599<br>751A - 516A 3.067<br>752A - 460A 4.595<br>752A - 516A 4.730<br>752A - 517A 3.270<br>752A - 518A 3.038<br>752A - 519A 4.080<br>755A - 513A 3.495<br>755A - 515A 3.484<br>755A - 516A 3.601<br>755A - 518A 4.441<br>756A - 519A 4.169<br>759A - 513A 2.296<br>759A - 520A 4.418<br>759A - 521A 3.464<br>759A - 522A 2.249<br>761A - 526A 3.193<br>762A - 526A 3.132<br>763A - 525A 3.014<br>763A - 526A 4.170 |

| | |
|---|---|
| | 375A - 12A 4.354<br>375A - 16A 4.784<br>376A - 280A 2.275<br>376A - 285A 4.173<br>376A - 287A 4.812<br>376A - 314A 4.423<br>379A - 285A 3.186<br>380A - 285A 2.546<br>383A - 285A 4.626<br>391A - 9A 3.138<br>391A - 12A 3.502<br>391A - 13A 3.057<br>393A - 4A 3.838<br>393A - 5A 4.044<br>393A - 6A 3.551<br>393A - 9A 3.797<br>394A - 6A 4.760<br>394A - 8A 4.488<br>397A - 3A 1.978<br>398A - 3A 3.804<br>399A - 3A 4.935<br>400A - 1A 2.801<br>400A - 3A 4.520<br>401A - 1A 2.770<br>401A - 2A 4.961<br>401A - 3A 2.604<br>682A - 42A 4.737<br>682A - 92A 2.429<br>684A - 92A 3.740<br>685A - 95A 1.608<br>703A - 42A 4.066<br>703A - 92A 4.145<br>720A - 45A 4.787<br>721A - 45A 3.612<br>722A - 45A 4.547<br>723A - 45A 2.818<br>725A - 45A 2.995<br>725A - 74A 2.847<br>725A - 75A 3.863<br>725A - 87A 4.388<br>725A - 89A 3.668<br>729A - 74A 3.109<br>729A - 75A 4.870<br>733A - 73A 4.062<br>740A - 141A 4.225<br>740A - 143A 4.705<br>747A - 144A 3.586 |

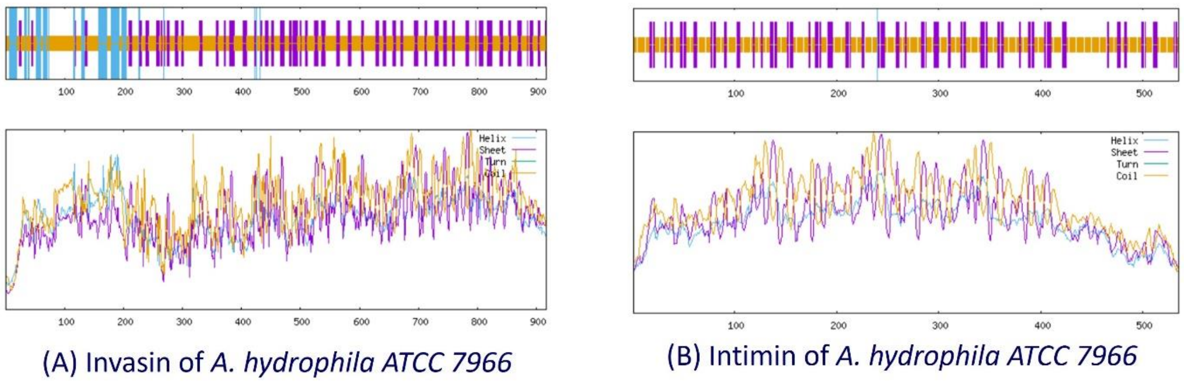

**Fig. S1:** Secondary structure analysis of (A) invasin and (B) intimin of *A. hydrophila* ATCC 7966 through SOPMA.

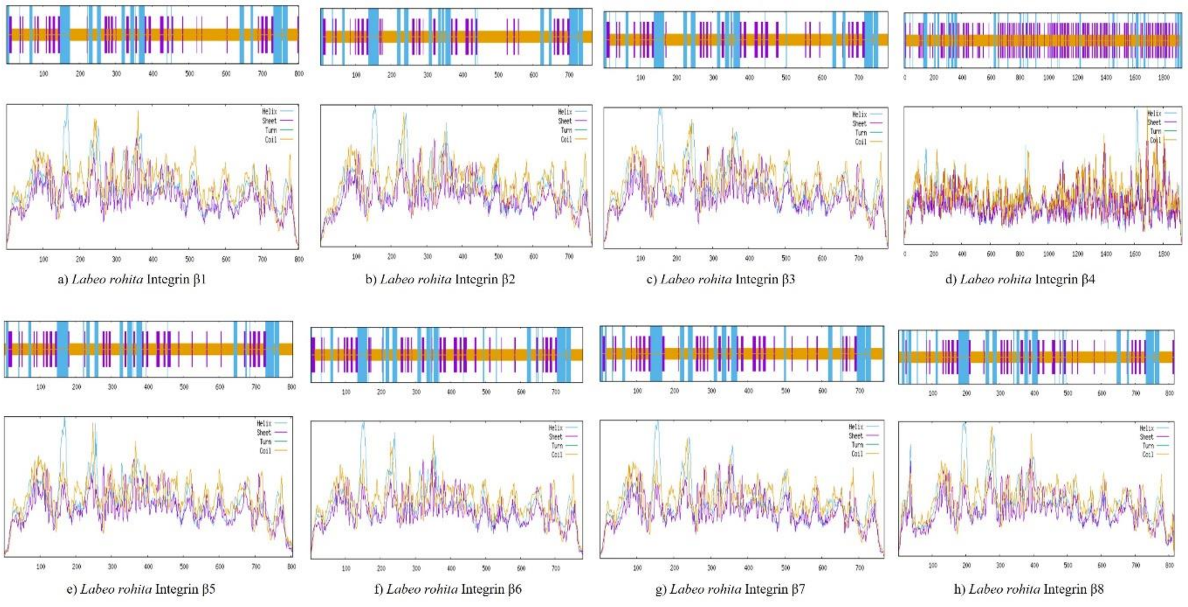

**Fig. S2:** Secondary structure analysis of β Intigrins -1 to -8 (a, b, c, d, e, f, g, and h) of *L. rohita* through SOPMA.

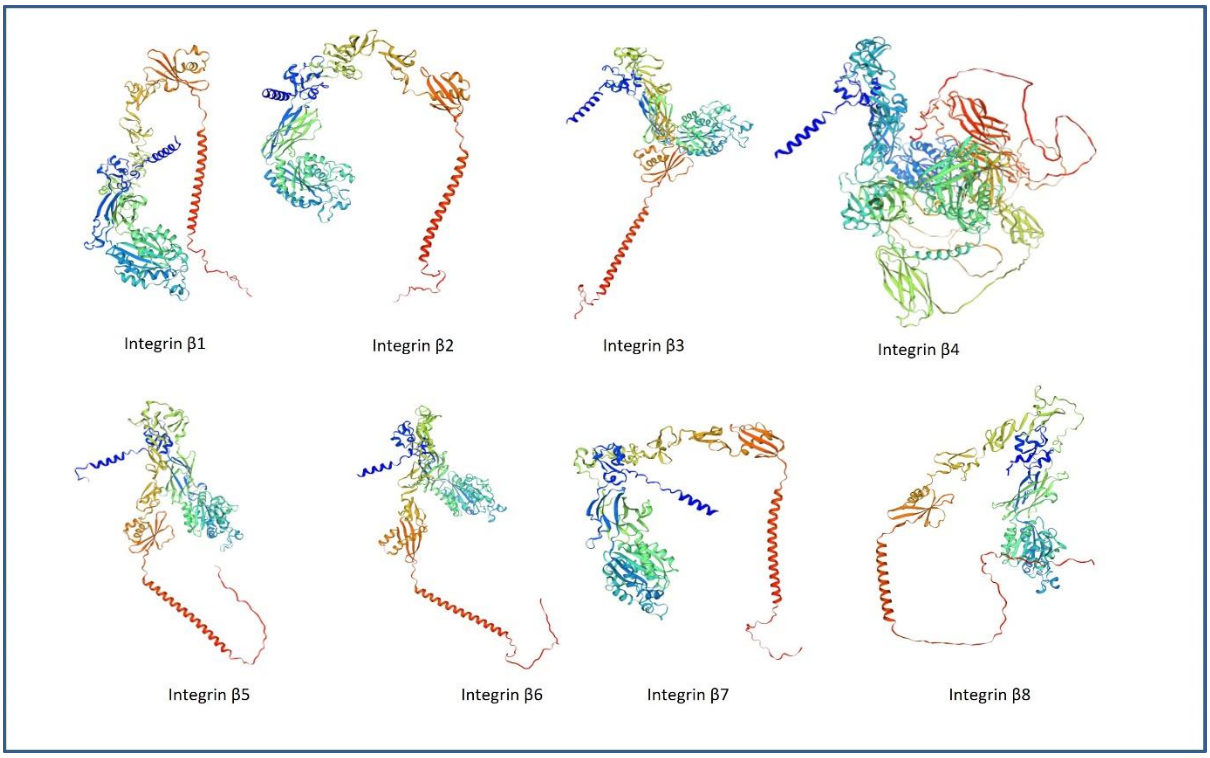

**Fig. S3:** Homology modeling of β Intigrins of *L. rohita* by SWISS-MODEL, accessed via the ExPASy web server.

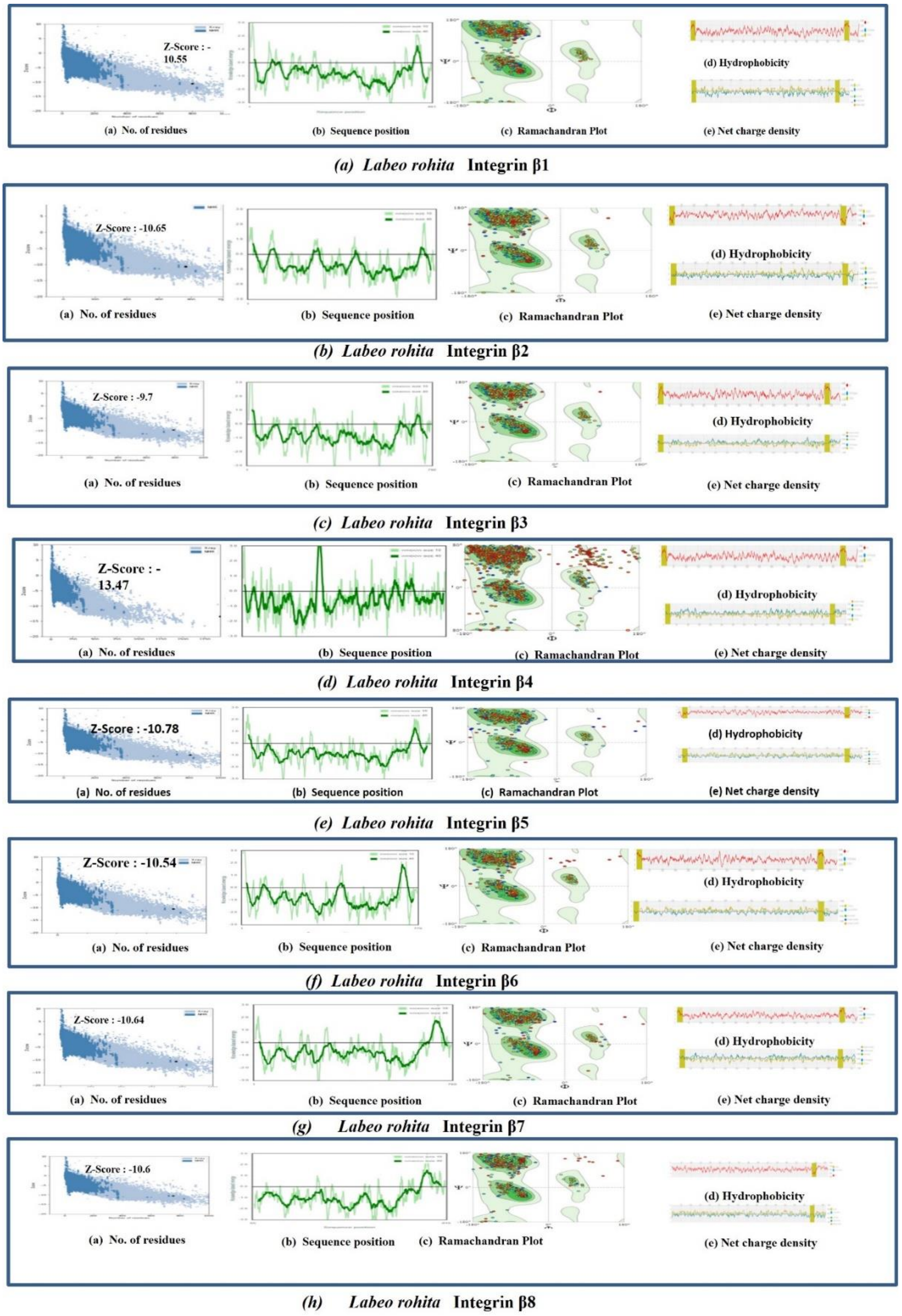

**Fig. S4:** Evaluation of tertiary structures of the β Intigrins -1 to -8 (a, b, c, d, e, f, g, and h) of *L. rohita*. In each panel, (a) represents Z- scores in terms of recediues, (b) represents sequence position, (c) represents Ramachandran plot, (d) represents hydrophobicity, and (e) represents net charge density.
